# Genetic interference of distinctive *Mycobacterium tuberculosis* peptidoglycan modifications enhances β-lactam susceptibility and reveals expression-sensitive host immune dynamics

**DOI:** 10.1101/2025.10.30.685640

**Authors:** Cátia Silveiro, Mariana Marques, Francisco Olivença, David Pires, Elsa Anes, Maria João Catalão

## Abstract

The high mortality associated with tuberculosis (TB), alongside the lack of efficient therapeutics against emerging multidrug-resistant *Mycobacterium tuberculosis* (*Mtb*) strains, emphasizes the need to identify novel antitubercular targets. Mycobacterial peptidoglycan, displaying characteristic modifications comprising the amidation of D-*iso*-glutamate (D-*i*Glu) and the *N*-glycolylation of muramic acid, is a promising therapeutic target. The genes encoding the enzymes mediating these PG modifications (*murT*/*gatD* and *namH*) were silenced in *Mtb* using CRISPR interference (CRISPRi) to investigate their impact on β-lactam susceptibility and host immune responses. First, qRT-PCR confirmed successful target mRNA knockdown, with variable repression efficiency based on the selected sgRNA, PAM strength, and target site. Phenotypic characterization through spotting dilution and growth curve assays corroborated the essentiality of D-*i*Glu amidation for mycobacterial survival, in contrast to the *N*-glycolylation of muramic acid. Moreover, susceptibility assays showed that both PG modifications contribute to β-lactam resistance, with sgRNA2-mediated *murT* knockdown substantially increasing β-lactam and isoniazid susceptibility. Furthermore, checkerboard assays showed reductions in the minimum fractional inhibitory concentration index (FICI_min_) value for AMX/MEM+CLA and EMB combinations following the depletion of both PG modifications, with significant differences observed upon *namH* knockdown. Additionally, D-*i*Glu amidation was uncovered as a determinant of *Mtb* survival within THP-1-derived macrophages at 6 days post-infection. Infection of THP-1-derived macrophages with MurT/GatD-depleted *Mtb* upregulated IL-1β and downregulated IL-10, whereas NamH depletion caused upregulation of both IL-1β and IL-10. Altogether, our findings unveiled the potential of targeting these PG modifications for the development of innovative therapeutic regimens against TB.

## INTRODUCTION

Despite improvements in diagnostics and chemotherapy, tuberculosis (TB) remains the deadliest infectious disease globally, causing over 1.25 million deaths and 10 million new cases annually (1). The spread of multidrug-resistant strains of *Mycobacterium tuberculosis* (*Mtb*), coupled with limited additions to the anti-TB drug pipeline, has exacerbated disease burden (2). The peculiar cell wall (CW) of *Mtb*, comprising a markedly modified peptidoglycan (PG), a profusely ramified arabinogalactan (AG), and long-chain mycolic acids (MA), is crucial for its antibiotic resistance and pathogenic success (3, 4). Since conventional TB chemotherapy targets AG and MA biosynthesis, novel antimycobacterials should exploit the unique subtleties of mycobacterial PG (mPG).

Among these, a distinctive feature of mPG is the amidation of the α-carboxyl group of D-*iso*-glutamate (D-*i*Glu) in both lipid I/II, catalyzed by the MurT/GatD amidotransferase complex (3, 4). GatD hydrolyses L-glutamine into glutamate and ammonia, which is channelled to the active site of MurT and incorporated into D-*i*Glu in an ATP/Mg^2+^-dependent fashion, generating D-*iso*-glutamine (D-*i*Gln) (5–8). Essential for mycobacterial survival, the *murT/gatD* operon is conserved among *Mtb* clinical isolates (8–10). The inhibition of D-*i*Glu amidation has been associated with impaired growth and increased β-lactam and lysozyme susceptibility in *Staphylococcus aureus* and *Mycobacterium smegmatis* (*Msm*) (9–13). Additionally, D-*i*Gln-containing lipid II is the favoured substrate for PG polymerization in *Mtb* (10, 14, 15) and may facilitate evasion of NOD1/2-mediated immune recognition (11, 16–18). Accordingly, mice vaccinated with a MurT/GatD-depleted *Mycobacterium bovis* Bacillus Calmette–Guérin (*M. bovis* BCG) strain exhibited heightened pro-inflammatory responses that controlled bacterial burden, granting superior protection versus BCG (19).

Another hallmark mPG modification is the *N*-glycolylation of muramic acid, catalyzed by the *N*-acetyl muramic acid hydroxylase (NamH), which hydroxylates the *N*-acetyl moiety to generate *N*-glycolyl (20). Consequently, mPG incorporates both *N*-acetyl and *N*-glycolyl muramic acid derivatives, interspersed with *N*-acetylglucosamine residues (20, 21). Although *namH* is non-essential for growth (22, 23), its strong conservation among *Mtb* clinical strains suggests it may play a part in mycobacterial fitness (9, 24). Notably, NamH depletion has been associated with increased β-lactam and lysozyme susceptibility (9, 20). Whereas some studies state that *N*-glycolylated PG stimulates NOD2-mediated immunity (25, 26), others suggest it may impair immune recognition (17).

Additional characteristic mPG features include the amidation of *meso*-diaminopimelic acid (*m*-DAP) on lipid II mediated by the AsnB amidotransferase and a preponderance of non-classical PG cross-links catalyzed by L,D-transpeptidases (Ldts), in detriment of 4→3 cross-links, catalyzed by penicillin binding proteins (PBPs) (27–29). The predominance of L,D-transpeptidation, resistant to conventional β-lactams (30), along with the impermeability of the MA layer, and the expression of a chromosomally encoded β-lactamase BlaC (31), have hindered the clinical use of β-lactams against *Mtb* (32). Nonetheless, carbapenem/β-lactamase inhibitor combinations (33) have shown efficacy against MDR-TB (34–39). Given the WHO’s call for further evidence on β-lactam efficacy in TB therapy, we have identified distinct β-lactam susceptibility patterns in *Mtb* (40, 41).

Recently, we demonstrated that the amidation of D-*i*Glu and the *N*-glycolylation of muramic acid modulate β-lactam resistance in *Msm* and contribute to its intracellular survival (9). To investigate the applicability of these findings in the pathogen, we implemented the CRISPRi platform (42) to target these PG modifications in *Mtb*.

## MATERIALS AND METHODS

### Bacterial strains, plasmids, culture conditions, cell lines, and antibiotics

The list of bacterial strains, plasmids, and cell lines used in this work are depicted in **Table 1**.

**Table 1.** Bacterial strains, plasmids, and cell lines used in this study.

| Bacterial strain, plasmid or cell line | Description | Reference or source |
| --- | --- | --- |
| <u>Bacteria</u> |  |  |
| *Escherichia coli* |  |  |
| JM109 | *recA1 endA1 gyr96 thi hsdR17 supE44 relA1 Δ(lac-proAB)*<br>[F' *traD36 proAB lac<sup>q</sup>ZΔM15*] | Stratagene |
| *Mycobacterium tuberculosis* |  |  |
| H37Rv | Virulent strain derived from *M. tuberculosis* H37 | 25618™, ATCC |
| <u>Plasmids</u> |  |  |
| PLJR965 (8631 bp) | Sth1 *dcas9* Tet<sup>R</sup> and Kan<sup>R</sup> L5 Int *attP* for *M. tuberculosis* | 42 |
| <u>Cell lines</u> |  |  |
| THP-1 | Human-derived monocyte cell line | TIB-202™, ATCC |

To amplify the CRISPRi backbone PLJR965 (Addgene ID: 115163) and for cloning, *Escherichia coli* (*E. coli*) strains were grown overnight (ON) at 37 °C with shaking in Luria-Bertani (LB) media (Merck). *Mtb* strains were grown in Middlebrook 7H9 broth (BD™ Biosciences) supplemented with 0.2% of glycerol (ThermoScientific), 10% OADC (oleic acid; albumin; dextrose; catalase), and 0.05% of tyloxapol (Sigma-Aldrich) or in 7H10 agar (BD™ Biosciences) plates supplemented with 0.5% of glycerol and 10% OADC at 37 °C, without shaking. The manipulation of *Mtb* strains was performed in a BSL3 laboratory. When appropriate, 25 μg/mL of kanamycin (NZYTech) were added to the media for plasmid selection. Target mRNA knockdown in mycobacterial cultures was induced by adding 100 ng*/*mL of ATc daily. To cultivate THP-1 cells, RPMI 1640 media (Gibco) was enriched with 10% of fetal bovine serum (Gibco), 1% of L-Glutamine (Gibco), 1% of HEPES (Gibco), 1% of sodium pyruvate (Gibco), and 1% of non-essential amino acids (Gibco). The cultures were maintained at 37 °C in the presence of 5% of CO_2_. Stocks of amoxicillin (AMX), cefotaxime (CTX), ethambutol (EMB), isoniazid (INH) and meropenem (MEM) (Sigma-Aldrich) were dissolved in sterile MilliQ water at a final concentration of 1.28 mg/mL (9). The ATc (Sigma-Aldrich) stocks were prepared at 10 mg/mL in dimethyl sulfoxide (DMSO) cell culture grade (AppliChem) (9). Potassium clavulanate (CLA) (Sigma-Aldrich) stocks were dissolved in phosphate buffer at 0.1 M, pH 6.0 (9).

### Construction of the knockdown mutants using CRISPRi

Cloning protocols were performed as previously detailed (9). To attain efficient knockdown, the coding strand or the promoter region of the genes of interest (GOIs) -*murT* (*Rv3712*), *gatD* (*Rv3713*), and *namH* (*Rv3818*) - were targeted. The strength of the chosen PAM is frequently correlated with the extent of knockdown efficiency and the ensuing growth inhibition phenotypes (42, 43). According to the predicted essentiality of the GOIs, their template strands were screened for PAM sequences (42). Strong PAMs were employed for the repression of putatively non-essential genes whereas medium-to-low strength PAMs mediated the knockdown of presumably essential genes. Besides accounting for gene essentiality, sgRNA design (**Table S1**) was optimized by combining distinct PAM strengths and target sites to achieve differential silencing profiles.

The CRISPRi backbone PLJR965 was amplified in *E. coli* JM109, digested, and purified as formerly explained (9). To clone the targeting sgRNAs, two complementary primers were designed (**Table S2**) and synthesized (Eurofins Genomics) with blunt ends to reconstitute the sgRNA promoter and the dCas9 handle sequences. This was accomplished following primer annealing and ligation of the annealed oligos to the backbone vector at 22 °C for 30 min, facilitated by T4 DNA ligase (400 U/µL; NEB).

To obtain the intended gene-targeting CRISPRi plasmids, chemically competent cells of *E. coli* were transformed with the ligation mixture by double heat-shock. Using the NZYMiniprep kit (NZYTech), recombinant DNA was isolated from the resultant transformants and then confirmed by Sanger sequencing to match the desired constructs. To generate the knockdown mutants (KDMs) in *Mtb*, electrocompetent cells were transformed with 300 ng of recombinant DNA in the presence of 10% glycerol, as previously reported (9, 44).

### Mycobacterial RNA isolation

To collect bacterial cultures for total RNA isolation, *Mtb* strains were grown to the exponential phase, normalized to a theoretical optical density at 600 nm (OD_600_) of 0.1-0.2, and divided into equal volumes devoid of or containing 100 ng/mL of ATc (43). After an incubation period of 72 h at 37 °C, approximately 6 x 10^8^ cells were harvested by centrifugation at 3,000 × *g* for 5-10 min at 4 °C (43), ressuspended and incubated in 500 μL of RNAprotect Bacteria Reagent (Qiagen) for 1 h, and stored at −80 °C. When necessary, the cells were thawed and pelleted by centrifugation at 5,000 × *g* for 5 min. To induce lysis, 350 µL of buffer NR (NZYTech) and 3.5 µL of β-mercaptoethanol (Merck) were added to the cells, which were then fully disrupted using BeadBug^TM^ (Benchmark Scientific). Bead-beating was carried out for 4 cycles at maximum speed, each lasting 1 minute, interleaved with 1-minute incubations on ice. After a spin-down, the lysate was loaded into the NZYSpin Homogenization column (NZYTech), and the workflow proceeded in conformity with the instructions provided by the NZYTotal RNA Isolation Kit. To elute RNA, the column was incubated with 20-30 μL of DEPC water (Invitrogen) at 60 °C for 10 min, followed by centrifugation and a second incubation-centrifugation step to maximize yield. A rigorous treatment with TurboDNase (Invitrogen) was then conducted to prevent contamination with genomic DNA. Nanodrop^TM^ was used to appraise the concentration and purity ratios of the eluted RNA and the efficacy of TurboDNase treatment was assessed by PCR amplification of the constitutive gene *sigA* (*Rv2703*), using NZYTaq DNA polymerase (NZYTech).

### Assessment of target gene repression by quantitative reverse-transcription PCR (qRT-PCR)

To confirm efficient repression of target genes, a qRT-PCR assay was performed. Briefly, 100 ng of purified RNA were reverse-transcribed into cDNA (9, 45) using the NZY First-Strand cDNA Synthesis Kit (NZYTech). Afterwards, a QuantStudio™ 7 Flex Real-Time PCR System (Applied Biosystems) was utilized to quantify cDNA following the preparation of qPCR reactions using the NZYSupreme qPCR Green Master Mix (2x), ROX (NZYTech). The qPCR program comprised an initial 2 min polymerase activation step at 95 °C, followed by 40 cycles of 5 s denaturation at 95 °C and 30 s annealing/extension at 60 °C. Using Primer 3 (https://primer3.ut.ee/), the primers were designed to produce a specific 100-200 bps amplicon (**Table S3**). By performing standard curves, the amplification efficiency of the primers was confirmed to be within the 80-110% range. Two technical replicates and a negative control were included in each run, and the average C_T_ was determined. Data analysis was performed using the Comparative C_T_ method, with *Mtb* WT serving as calibrator and *sigA* as the internal control gene (42, 43, 46). The quantification of target mRNA sequences was performed using three biological replicates.

### Phenotyping assays

To ascertain the importance of distinctive mPG modifications to bacterial growth, phenotypic characterization was performed using spotting dilution and growth curve assays.

Spotting dilution assays were conducted as previously described (9). Briefly, *Mtb* cultures were grown to the exponential phase, adjusted to a theoretical OD_600_ of 0.001, and subjected to 1:2 dilution in a 96-well plate. Then, 5 μL of each bacterial suspension were plated onto 7H10 + OADC (+ATc) plates at 37 °C for approximately 15 days. More than three independent experiments were completed to ensure assay reproducibility.

To monitor bacterial growth, *Mtb* liquid cultures were grown to the exponential phase, normalized to a theoretical OD_600_ of 0.005-0.01 and, further incubated at 37 °C with and without inducer for 264 hours. The concentration of ATc was replenished to 100 ng/mL every 24 h. The OD_600_ was measured at 96, 120, 168, 192, and 164 h across three independent experiments. Growth was evaluated by plotting the mean OD_600_ values versus the incubation time for each strain under study.

### Minimum inhibitory concentration (MIC) determination

To unveil changes in antibiotic susceptibility upon target gene repression, the minimum inhibitory concentrations of three β-lactams (amoxicillin, cefotaxime, and meropenem), in the presence and absence of the β-lactamase inhibitor clavulanate, and of two first-line antimycobacterials (isoniazid and ethambutol) were ascertained. Succinctly, the compounds under assessment were diluted to their predefined initial concentrations (512 μg/mL for β-lactams and 128 μg/mL for antitubercular agents), whereas clavulanate was used at a final concentration of 2.5 μg/mL. The antimicrobials were then subjected to a 1:2 serial dilution in supplemented 7H9 media. In the last two rows, positive (bacteria only) and negative (media only) control wells were included. The cultures of *Mtb* were first diluted to a theoretical OD_600_ of 0.1 and split into two equal volumes, one of which was subjected to a pre-depletion period of 3 days with 100 ng/mL of ATc (47). Following this period, the cultures were normalized to a OD_600_ of 0.002, and the bacteria were induced with a final concentration of 100 ng/mL of ATc. After, 100 μL of every bacterial suspension were dispensed into the designated wells to a final concentration of 10⁵ CFU/mL, and the plates were then incubated at 37 °C for 10-12 days. The MIC was annotated as the minimum concentration of antibiotic that completely suppressed observable bacterial growth. Three independent experiments were carried out for each strain, and the median MIC values were calculated.

### Disk diffusion assays

To assess potential changes in antibiotic susceptibility on solid media upon induced target interference, disk diffusion assays were performed with AMX+CLA, CTX+CLA, and MEM+CLA, following previously published protocols (31). Briefly, *Mtb* cultures were grown to the exponential phase and diluted to a theoretical OD_600_ of 0.1. To create a mycobacterial lawn, bacterial suspensions were then transferred and spread onto the 7H10 + OADC (+ATc) plates, which were allowed to dry for 20 min, and incubated at 37 °C for 1-2 days. Afterwards, β-lactams were diluted to the required concentrations for susceptibility testing (AMX: 512 µg/mL; CTX: 512 µg/mL; MEM: 128 µg/mL; CLA: 5 µg/mL), and 20 μL of each compound were transferred to the respective 9 mm filter paper disks (Filterlab), which were allowed to dry and placed onto the mycobacterial lawn using a pipette tip. Following 10–13 days of incubation at 37 °C, the diameter of the zone of inhibition (halo) was measured using a ruler. The results depict the mean diameter of the zone of inhibition (in cm) from a minimum of three biological replicates.

### Checkerboard assays

To evaluate the interactions between frontline agents (isoniazid or ethambutol), β-lactams, and the CRISPRi-mediated interference of target genes (simulating the prospective chemical inhibition of NamH and MurT/GatD), checkerboard assays were performed. Since previous observations showed a synergistic effect between EMB and amoxicillin-clavulanate (AMX+CLA) or meropenem-clavulanate (MEM+CLA) in *Mtb* H37Rv (48), we selected these combinations for further antibiotic interaction assessment. The minimum fractional inhibitory concentration indices (FICIs) were calculated to measure the extent of these interactions and whether these were classified as synergy (FICI ≤ 0.5), additive effect (0.5 < FICI ≤ 1), indifference (1 < FICI ≤ 4), or antagonism (FICI > 4) (49, 50).

The checkerboard assays were conducted as previously described (48), with a few modifications. Two or three days before the assay, *Mtb* cultures were diluted to a theoretical OD_600_ of 0.1 and split into two equal volumes, one of which was subjected to a pre-depletion period of 2-3 days with 100 ng/mL of ATc (47). On the day of the assay, the antibiotics were prepared at four times (dilution factor) the initial test concentration (corresponding to the quadruple of the respective individual MIC) and serially diluted 1:2 in supplemented 7H9. Using a multichannel pipette, 50 µL of each antibiotic under study was transferred to the wells of a 96-well plate, along the abscissa or ordinate, thus forming an 8 x 8 matrix with a total volume of 100 µL. To preserve a humidified atmosphere, water or medium was added to the remaining wells. The cultures were then centrifuged at 3,000 × *g* for 5 min, resuspended in 1 mL of supplemented 7H9 media, submitted to an ultrasonic bath for 5 min, and centrifuged at 500 × *g* for 15 s to obtain a clump-free suspension. After, the cultures were diluted to a theoretical OD_600_ of 0.002 in supplemented 7H9 media, to which clavulanate (5 μg/mL) and ATc (100 ng/mL) were added at the indicated final concentration, when appropriate. After, the matrix wells were inoculated with 100 µL of each bacterial suspension, producing a final concentration of 10⁵ CFU/mL. The OD₆₀₀ of the wells was measured using an Infinite M200 Pro microplate reader (TECAN) following 10–12 days of incubation at 37 °C.

In this case, the MIC was annotated as the concentration achieving ≥ 99% inhibition of mycobacterial growth relative to the positive control. The fractional inhibitory concentration (FIC) of each tested antibiotic was determined by dividing the MIC of the antibiotic when used in combination by its individual MIC (e.g., FIC_EMB_ = MIC_EMBxAMX+CLA_/MIC_EMB_). The FIC index (FICI) is obtained by adding the individual FICs of the antibiotics being tested in each well (e.g., FICI for interaction assessment between EMB and AMX+CLA = MIC_EMBxAMX+CLA_/MIC_EMB_ + MIC_EMBxAMX+CLA_/MIC_AMX+CLA_). The presented results show the FICI values as the mean ± SEM of at least three independent experiments.

### THP-1 infection assays

To investigate the role of the distinctive modifications of mPG in the intracellular survival of *Mtb*, THP-1-derived macrophages were infected with the control and mutant strains. THP-1 monocytes were routinely passaged every 3–4 days to maintain a density of 2 × 10⁵ cells/mL. As needed, the cells were seeded at a density of 1.5 × 10⁵ cells/well in 48-well flat-bottom tissue culture plates and treated with 20 nM phorbol 12-myristate 13-acetate (PMA; Sigma-Aldrich) for 72 hours to induce differentiation into resting macrophages (48, 51, 52). On the day of THP-1 seeding, *Mtb* cultures were diluted to an OD_600_ of approximately 0.05 and split into uninduced and induced conditions, which were treated with 100 ng/mL of ATc (47). The cultures were then incubated at 37 °C for 72 h, after which the concentration of inducer was replenished.

One day prior to infection, the cell media containing PMA was exchanged by prewarmed supplemented PMA-deprived RPMI media. For the infection, log phase *Mtb* cultures were harvested by centrifugation at 3,000 × *g* for 5 min, washed with 5 mL of phosphate-buffered saline (PBS) 1 X (Gibco), and resuspended in supplemented RPMI media (48, 51, 53–55). The bacterial suspension was then submitted to an ultrasonic bath for 5 min, and the remaining clumps were pelleted by centrifugation at 500 × *g* for 15 s, enabling the recovery of a single-cell suspension (48, 51, 53–56). Following the removal of the cell media from the plates, THP-1-derived macrophages were infected with the mycobacterial suspensions at a multiplicity of infection (MOI; ratio of bacteria/cells) of 0.25 and, incubated at 37 °C with 5% CO_2_ for various timepoints (1 d; 3 d; 6 d), consistent with a long-term infection model that minimizes macrophage cell death (57, 58). At this point, induced cultures were added 100 ng/mL of ATc. Following an internalization period of 3 h, the macrophages were washed twice with PBS 1 X to remove extracellular bacteria and cultivated in prewarmed supplemented RPMI media (48, 51, 54), devoid of or containing 250 ng/mL of ATc, necessary to sustain long-term target knockdown (47). Prewarmed supplemented RPMI media and ATc (225 ng/mL) were replenished 3 days after infection onset (T_3_) to sustain cell growth (47).

### Intracellular survival evaluation

After the specified incubation times (T_1_, T_3_, and T_6_), the medium was discarded, and cell lysis was promoted by incubating the infected macrophages with a 0.05% IGEPAL solution (Merck) at 37 °C for 15 min (48, 54, 56). Subsequently, the lysates were serially diluted in sterile water and plated onto 7H10 + OADC plates, which underwent incubation at 37 °C with 5% CO_2_ for 2-3 weeks until colonies were visible (48, 54, 56). Finally, the colony-forming units (CFU) from three independent biological replicates, each comprising three technical replicates, were counted using an Olympus CK2 Inverted Phase Contrast Microscope.

### Immune response assessment of infected THP-1-derived macrophages via qRT-PCR

To ascertain whether the amidation of D-*i*Glu or the *N*-glycolylation of muramic acid induce differential immune responses upon infection with *Mtb*, THP-1-derived macrophages were infected with each of the study strains. After 6 days of infection, THP-1 cells were lysed and transcript levels of pro-inflammatory cytokines tumor necrosis factor alpha (TNF-α) and interleukin-1 β (IL-1β), and of the anti-inflammatory cytokine interleukin-10 (IL-10), were measured by qRT-PCR. For these assays, THP-1 monocytes were seeded at a density of 5 × 10⁵ cells/well in 12-well flat-bottom tissue culture plates and treated with 20 nM phorbol 12-myristate 13-acetate (PMA; Sigma-Aldrich) for 72 h to induce differentiation into resting macrophages (48, 51). The preparation of bacterial cultures and macrophage infection (MOI of 0.25) proceeded as aforementioned. The lysis of infected macrophages was promoted by vigorous cell resuspension in 394 µL of buffer NR (NZYTech) and 4 µL of β-mercaptoethanol (Merck). The obtained lysates were transferred to an NZYSpin Homogenization column (NZYTech), and the ensuing steps were performed following the instructions of the NZYTotal RNA Isolation Kit (55, 56). The isolation of purified RNA was facilitated by a two-step elution with 20-30 μL of DEPC-treated water (Invitrogen) at 60 °C, and the quantity and integrity of the isolated RNA were evaluated using Nanodrop™. Then, 100 ng of total RNA were used as template for cDNA synthesis using the NZY First-Strand cDNA Synthesis Kit (NZYTech) (56). The qRT-PCR assay was carried out as previously described, with the following primers, designed to amplify a specific 100-200 bps amplicon and possessing an amplification efficiency within the 80-110% range (**Table S4**). Each run included two technical replicates and a negative control, and the average C_T_ value was calculated. Data analysis was carried out using the Comparative C_T_ method, with non-infected macrophages as the calibrator sample and *GAPDH* (*glyceraldehyde 3-phosphate dehydrogenase*) as the endogenous control gene (51, 55, 56). The mRNA levels of TNF-α, IL-1β, and IL-10 were quantified using three biological replicates.

### Statistical analysis

All statistical analyses were carried out using GraphPad (version 8.4.3.686), and results are expressed as mean ± SEM, unless mentioned otherwise. The Shapiro-Wilk test was employed in normality testing. The ordinary one-way ANOVA was used to compare multiple groups, followed by pairwise analyses of preselected groups using the Holm-Sidak test. All the prerequisites for the employed tests were evaluated and confirmed.

## RESULTS

### Construction of CRISPRi knockdown mutants

To engineer the knockdown mutants of our GOIs in *Mtb*, the ATc-regulated dCas9_Sth1_-mediated CRISPRi system was employed (42). Distinct PAM sequences were identified in the template strand of the GOIs to assess differential knockdown efficiency owing to PAM strength and/or target site (9) (**Figure 1A**). To prevent lethal phenotypes, PAMs mediating target interference were selected based on the predicted essentiality of the GOIs (9, 52). Since our GOIs are operonic, the local gene context was also considered to deter polar effects (42, 45, 46) (**Figure 1B**). The interference plasmids were verified by Sanger Sequencing (**Figure S1**).

**Figure 1.**
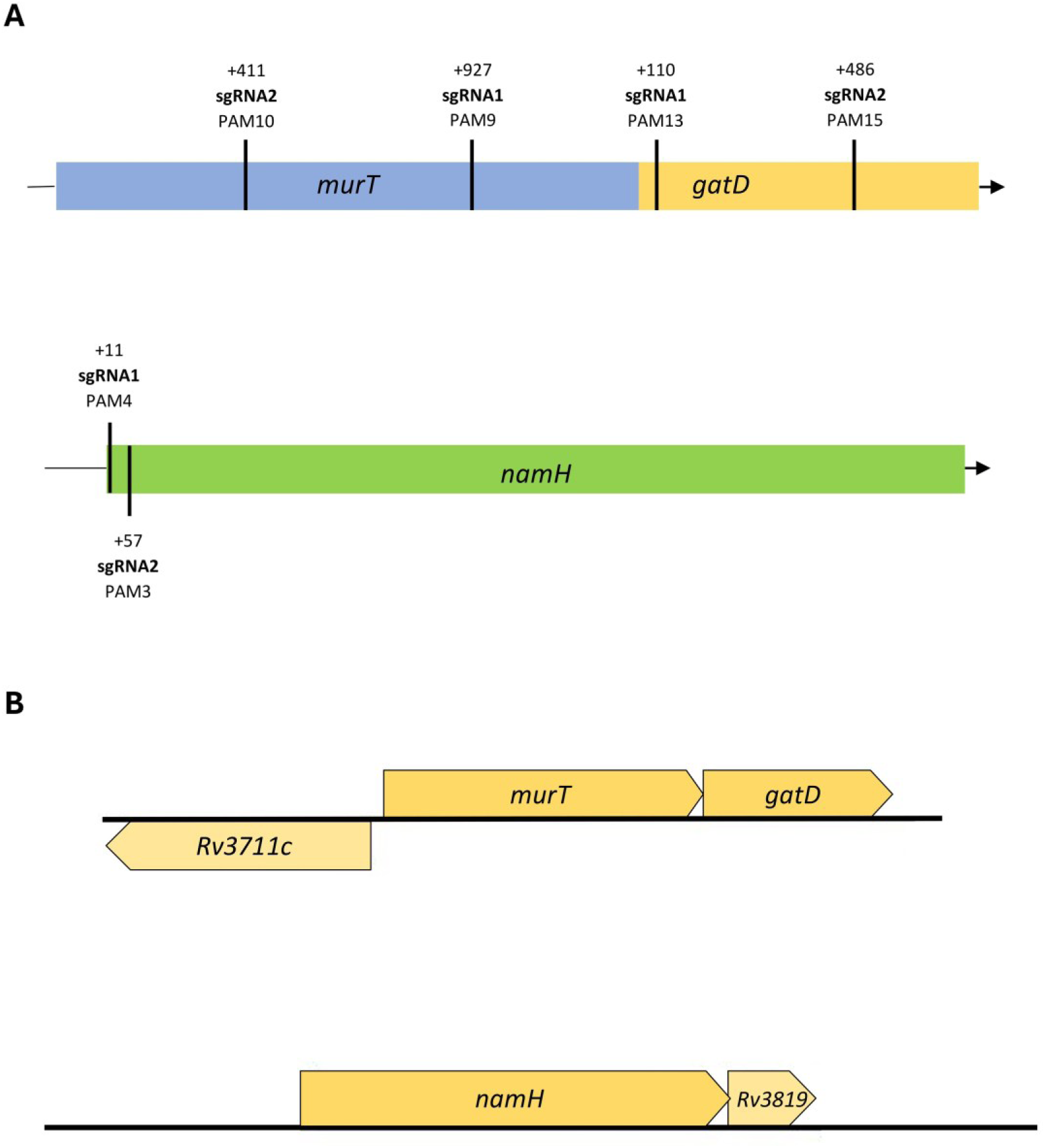
(A) Distribution of sgRNA targeting sequences along the *murT*/*gatD* operon and the *namH* gene in *Mtb*. Different sites were targeted along the genes to assess the differential repression efficiency of CRISPRi. (B) The gene local context of the target genes *murT*, *gatD*, and *namH* should be considered due to potential downstream/upstream polar effects. The *murT* and *gatD* genes are co-transcribed under regulation of a single promoter. The *Rv3711c* gene precedes *murT* but is transcribed in the opposite direction. A single promoter guides the conjoint transcription of the *namH* and *Rv3819* genes by RNA polymerase.

The normalized mRNA expression of *murT*, *gatD, namH,* and their adjacent genes was determined in the control strains *Mtb* WT and PLJR965, and in the KDMs, with and without inducer, at 72 h post-induction (**Figure 2A,B**).

**Figure 2.**
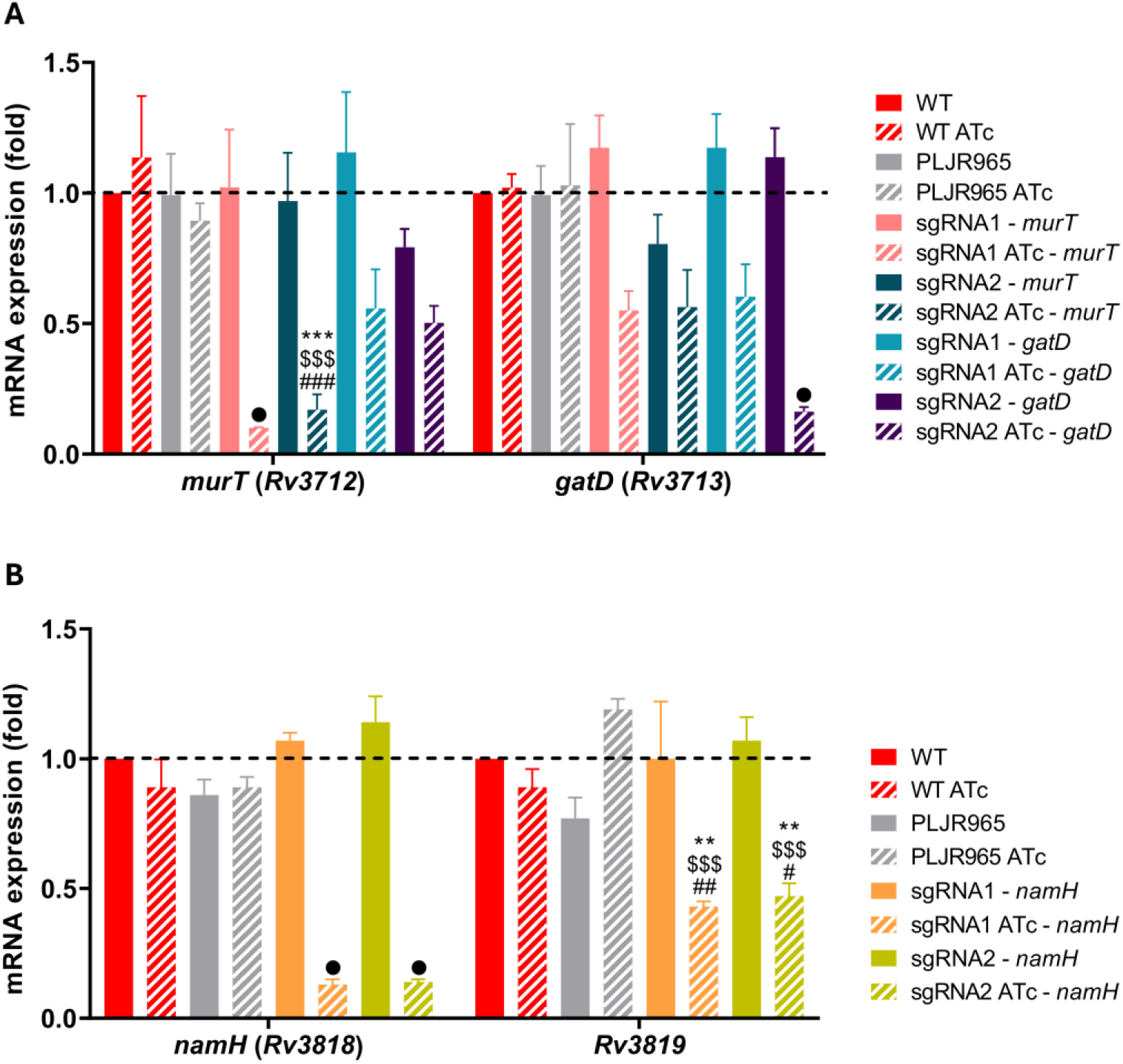
Mean relative mRNA expression levels of target genes, normalized to *sigA*, at 72 hours post-induction, with (stripped bars) and without (smooth bars) 100 ng/mL ATc (n=3): (A) *murT* and *gatD* knockdown; (B) *namH* knockdown. The dashed lines show the WT sample as calibrator. Error bars show the standard error of the mean (SEM). Multiple comparisons were made using one-way ANOVA: * *P* < 0.05; ** *P* < 0.01; *** *P* < 0.001. Significant differences are indicated with symbols: # compared to *Mtb* WT ATc, $ compared to *Mtb* PLJR965 ATc, * versus the respective uninduced control. The • symbol indicates a highly significant difference (*P* < 0.0001) from all controls.

The results demonstrate that the *murT* gene was successfully silenced in all induced KDMs (**Figure 2A**). The sgRNA1-directed silencing of *murT* caused a highly significant reduction in its transcript levels (*P* < 0.0001) relative to WT ATc (11.3-fold), PLJR965 ATc (8.9-fold), and the respective uninduced control (10.1-fold). Likewise, sgRNA2-directed *murT* knockdown caused a very significant repression versus WT ATc (6.7-fold; *P* = 0.0002), PLJR965 ATc (5.2-fold; *P* = 0.0005), and the respective uninduced control (5.7-fold; *P* = 0.0005). Differences in PAM strength and sgRNA length directly accounted for the greater repression of *murT* attained with sgRNA1-*murT* (PAM9, +927 bp) compared to sgRNA2-*murT* (PAM10, +411 bp). In contrast, only one of the targeting sgRNAs efficiently repressed *gatD* (**Figure 2A**). Indeed, sgRNA2-mediated *gatD* knockdown caused a highly significant decrease in its expression (*P* < 0.0001) versus WT ATc (6.2-fold), PLJR965 ATc (6.3-fold), and the respective uninduced control (7.0-fold). No significant polar effects were observed in adjacent genes upon CRISPRi-mediated silencing of the *murT*/*gatD* operon (**Figure S2**).

Additionally, the *namH* gene was successfully silenced in all induced KDMs (**Figure 2B**). A highly significant knockdown (*P* < 0.0001) was attained with both sgRNA1-*namH* and sgRNA2-*namH* compared to WT ATc (6.9-fold *vs*. 6.4-fold), PLJR965 ATc (6.9-fold *vs*. 6.4-fold), and the respective uninduced mutant (8.3-fold). The repression efficiency of sgRNA1-*namH* (PAM4, +11 bp) and sgRNA2-*namH* (PAM3, +57 bp) is comparable, as repression differences between strong PAMs are negligible (42). Moreover, both sgRNAs target the 5’ region of *namH*: sgRNA1-*namH* targets the promoter region, while sgRNA2-*namH* targets the first nucleotides of the ORF. Both sgRNAs caused significant downstream polar effects on *Rv3819* (**Figure 2B**), which are likely to partially affect the ensuing phenotypes. While non-essential (22, 23, 59, 60) and functionally uncharacterized, *Rv3819* encodes a protein containing a PS00012 phosphopantetheine attachment site.

### Phenotyping reveals the essentiality of *murT*/*gatD* and non-essentiality of *namH*

To phenotypically characterize the KDMs, spotting dilution and growth curve assays were conducted (**Figures 3 and 4**).

**Figure 3.**
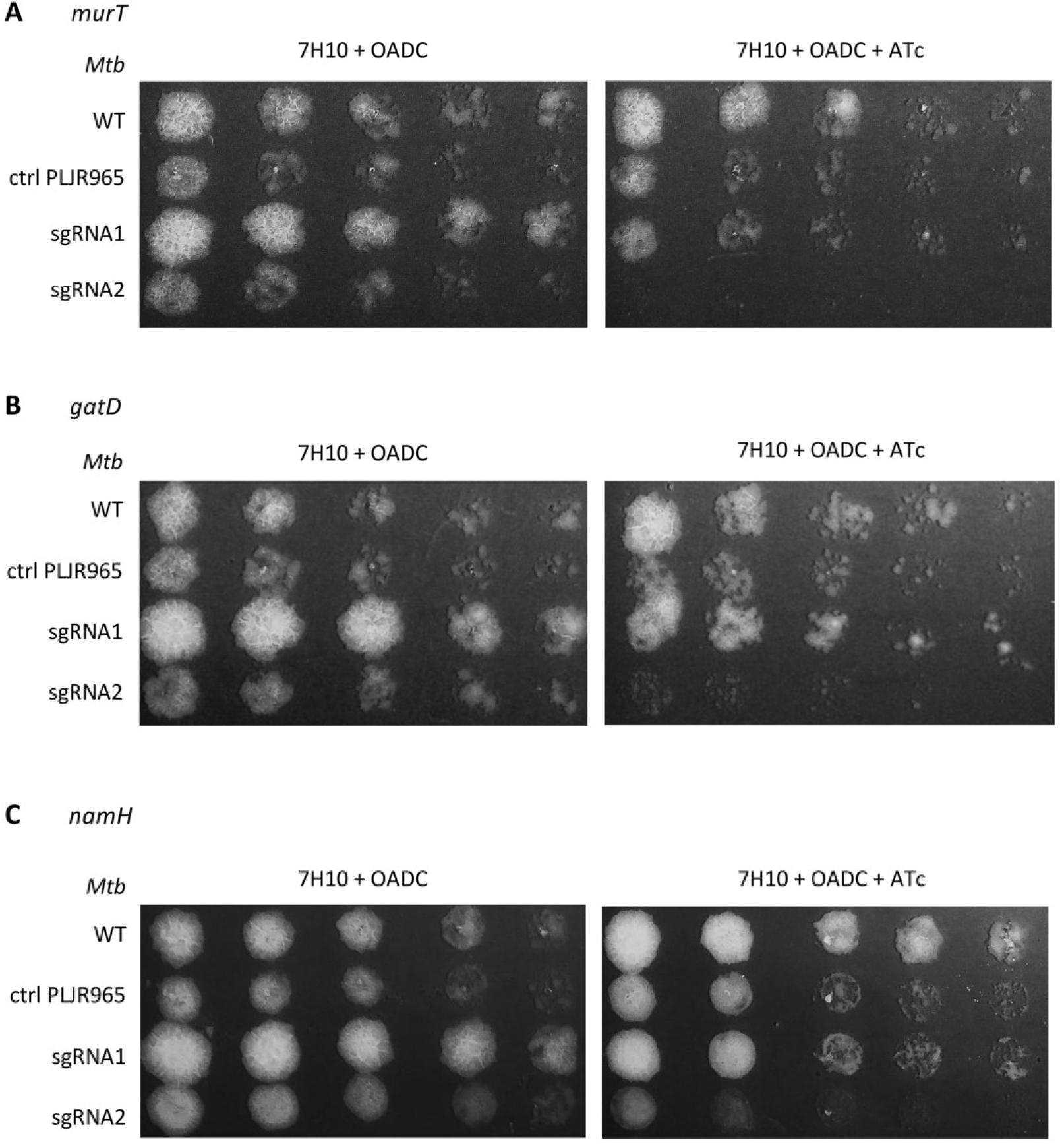
Spotting dilution assays of the *M. tuberculosis* knockdown mutants (n=3): (A) *murT* knockdown mutants; (B) *gatD* knockdown mutants; (C) *namH* knockdown mutants. Negative controls (*Mtb* WT and PLJR965) were included in all experiments. Each culture was first normalized to an OD600 of 0.001, then serially diluted 2-fold, and spotted onto agar plates.

**Figure 4.**
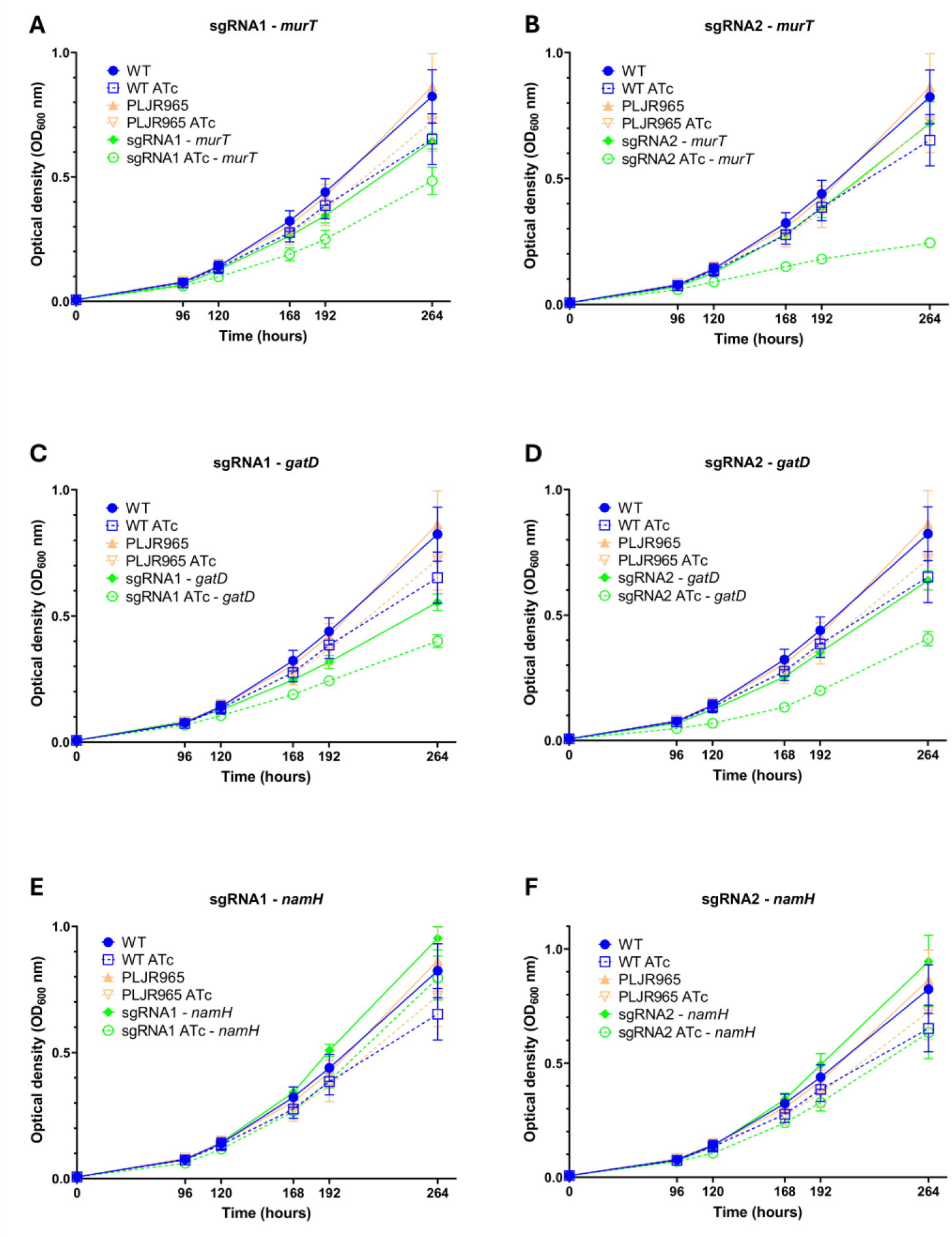
Growth curves of the *M. tuberculosis* knockdown mutants, in the absence or presence of 100 ng/mL of ATc (n=3): (A, B) *murT* knockdown mutants; (C, D) *gatD* knockdown mutants; (E, F) *namH* knockdown mutants. Negative controls (*Mtb* WT and PLJR965) were included in all experiments. Error bars show the standard error of the mean (SEM).

The spotting dilution assays demonstrated that the growth of the control strain *Mtb* WT was not impacted by inducer addition. In contrast, *Mtb* PLJR965 displayed only a minimal growth reduction on solid media containing ATc, suggesting that the expression of exogenous dCas9_Sth1_ may be somewhat toxic to *Mtb*. The results confirmed the essentiality of the *murT*/*gatD* operon for *Mtb* survival, as its knockdown caused severe growth impairment, particularly when mediated by potent sgRNAs. In the case of *murT* knockdown (*murT*^-^), sgRNA2-mediated silencing (PAM10, +411 bp) completely abrogated *Mtb* growth, while sgRNA1-*murT* (PAM9, +927 bp) provoked only a modest growth defect (**Figure 3A**). Although the employed sgRNAs theoretically mediate comparable levels of target repression, sgRNA2-*murT* guides dCas9_Sth1_ binding to the 5’-region of *murT*, causing RNA polymerase dissociation at an early stage of transcription, and resulting in the production of a small mRNA likely to be degraded. On the other hand, sgRNA1-*murT* targets a site near the 3’-region of *murT*, with dCas9_Sth1_-mediated steric blockage of the RNA polymerase leading to the production of a larger mRNA, likely to produce a semi-functional MurT protein. Regarding *gatD* knockdown (*gatD^-^*), sgRNA1-*gatD* (PAM13, +110 bp) caused a modest growth reduction whereas sgRNA2-*gatD* (PAM15, +486 bp) severely impacted bacterial survival on solid media, despite employing a theoretically “weaker” PAM (**Figure 3B**). The observed phenotypes reflect the greater knockdown mediated by sgRNA2-*gatD* compared to sgRNA1-*gatD*. Moreover, sgRNA1-mediated *namH* repression caused a noticeable but modest reduction in colony size, whereas sgRNA2-mediated *namH* knockdown provoked an observable growth deficit (**Figure 3C**). These observations suggest that the activity of NamH impacts the growth of *Mtb* in these conditions.

The growth curves showed that *murT* repression severely affected bacterial growth in liquid media (**Figure 4A,B**). Specifically, sgRNA2-mediated silencing (PAM10, +411 bp) led to a severe reduction in the growth rate of *Mtb*, while sgRNA1-*murT* (PAM9, +927 bp) caused only a modest growth defect. These observations are in line with the spotting dilution assays (**Figure 3A**). Regarding *gatD* repression (*gatD*^-^), sgRNA2-mediated knockdown caused a more substantial decrease in the growth rate of *Mtb* (T_120_, T_168_, and T_192_ h), compared to sgRNA1-*gatD* (**Figure 4C,D**). This observation is expected, since *gatD* silencing is more efficient with sgRNA2 (**Figure 2A**) and in accordance with the spotting assays (**Figure 3B**). As for *namH* repression (*namH*^-^), sgRNA1-*namH* did not provoke any observable changes in *Mtb* growth while sgRNA2-*namH* provoked a subtle growth reduction (**Figure 4E,F**), consistent with the spotting dilution assays (**Figure 3C**).

### The activity of MurT/GatD and NamH contributes to β-lactam resistance

To determine how the characteristic modifications of mPG influence the antibiotic susceptibility profile of *Mtb*, the MICs of ethambutol, isoniazid, and three β-lactams, in the absence or presence of clavulanate, were determined against the control strains and the KDMs (**Figure 5A**). As predicted, ATc induction did not considerably affect the MIC values for the control strains (*Mtb* H37Rv WT, *Mtb*::PLJR965). Conversely, all KDMs displayed increased susceptibility to the tested β-lactams, irrespective of the targeted gene.

**Figure 5.**
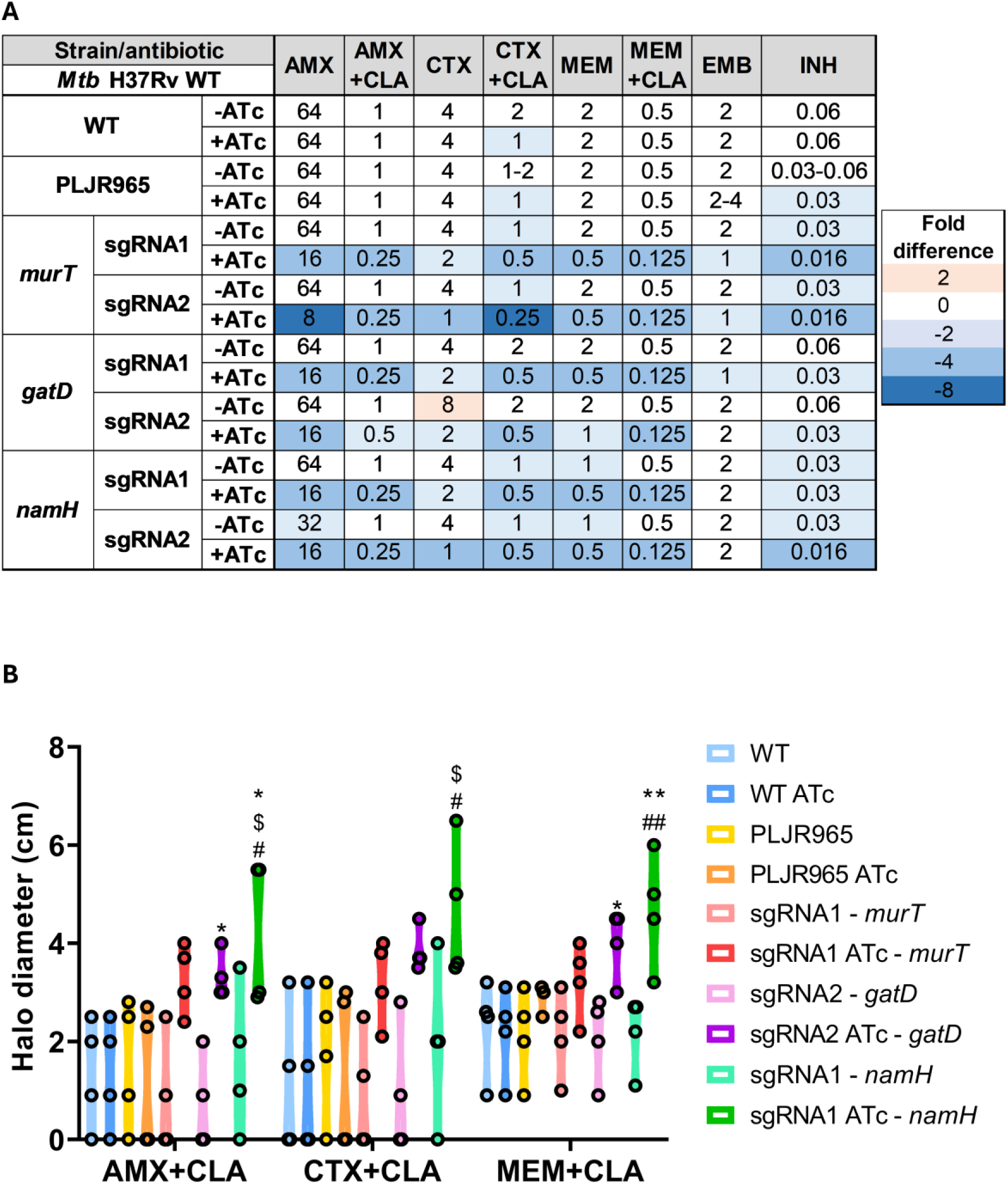
(A) Heatmap of the fold differences in the minimum inhibitory concentrations (MICs) of the knockdown mutants constructed in *M. tuberculosis*, with and without 100 ng/mL of ATc. Columns represent the antibiotics and rows represent the strains. The median MIC (in μg/mL) of each antibiotic for each strain is shown in the respective intersection, and colours depict an increase (red shades) or decrease (blue shades) in the value when compared to the MICs of *Mtb* WT. AMX, amoxicillin; CLA, clavulanate; CTX, cefotaxime; EMB, ethambutol; INH, isoniazid; MEM, meropenem. (B) Violin plot of the diameter of the zone of inhibition (in cm) for each combination of beta-lactam/CLA with the indicated knockdown mutants constructed in *Mtb* WT for the *murT* (sgRNA1), *gatD* (sgRNA2) and *namH* (sgRNA1) genes with and without 100 ng/mL of ATc (n ≥ 3). AMX, amoxicillin; CLA, clavulanate; CTX, cefotaxime; MEM, meropenem. Violin plots were clipped at y = 0, as negative halo diameters are not biologically feasible and open black dots represent individual data points. Multiple comparisons were made using one-way ANOVA, with significance levels: * *P* < 0.05; ** *P* < 0.01; *** *P* < 0.001. Significant differences are indicated with symbols: # comparing to *Mtb* WT ATc, $ comparing to *Mtb* PLJR965 ATc, * comparing to the respective uninduced control.

Generally, the median MICs for the KDMs revealed that depletion of the amidation of D-*i*Glu (*murT*/*gatD*^-^) and of the *N*-glycolylation of muramic acid (*namH*^-^) promoted increased susceptibility to β-lactams. Regarding *murT* and *namH* repression, the increased β-lactam susceptibility phenotypes were well correlated with the growth deficits displayed by the respective KDMs in the spotting dilutions (**Figure 3 A,C**). Despite sgRNA1-*murT* being more efficient at silencing *murT* (**Figure 2A**), sgRNA2-mediated *murT* knockdown resulted in a lethal phenotype in the spotting dilution assays (**Figure 3A**) and heightened β-lactam susceptibility (**Figure 5A**). Besides, sgRNA2-mediated *namH* silencing not only caused a growth reduction in the spotting dilution assays (**Figure 3C**) but also provoked a β-lactam hypersusceptibility phenotype (**Figure 5A**). Surprisingly, the sgRNA1-mediated silencing of *gatD* did not result in a significant knockdown (**Figure 2B**) but caused greater β-lactam susceptibility differences than sgRNA2-mediated *gatD* silencing (**Figure 5A**). It seems that a minimal repression mediated by a sgRNA that targets the 5’-region of *gatD* has a greater impact on β-lactam susceptibility than significant repression mediated by a sgRNA targeting a region adjacent to the 3’-end of *gatD*. Overall, sgRNA2-mediated *murT* silencing caused the largest MIC differences compared to *Mtb* WT, with the MICs of AMX and CTX+CLA decreasing by 8-fold (**Figure 5A**).

For AMX, all induced KDMs displayed substantial susceptibility differences (≥ 4-fold) relative to *Mtb* WT. Likewise, considerable susceptibility differences in AMX+CLA were observed in all KDMs except for the induced *Mtb*::PLJR965-sgRNA2-*gatD* KDM. In fact, the MICs of CTX remained mostly constant, except for the induced *Mtb*::PLJR965-sgRNA2-*murT* and *Mtb*::PLJR965-sgRNA2-*namH* KDMs, which exhibited a striking increase in β-lactam susceptibility. Regarding CTX+CLA, all induced KDMs showed considerable susceptibility differences (≥ 4-fold) relative to *Mtb* WT. Regarding MEM, substantial susceptibility differences (≥ 4-fold) were found in all KDMs except for the induced *Mtb*::PLJR965-sgRNA2-*gatD* KDM. In the case of MEM+CLA, all induced KDMs presented considerable susceptibility differences versus *Mtb* WT. The addition of CLA to β-lactams provoked a notable 64-fold reduction in the MICs of AMX, while the MICs of CTX and MEM were only reduced by 4-fold. The MIC of MEM+CLA against the induced KDMs was 0.125 μg/mL, being 4-fold lower than the WT MIC. Altogether, MEM+CLA seems to synergize well with MurT/GatD and NamH inhibition. Among the tested β-lactams, MEM consistently exhibited the lowest MICs, being the most effective at inhibiting *Mtb* growth. Finally, the MICs of EMB remained stable, while three induced KDMs displayed increased susceptibility to INH, with the effect being most pronounced in the *murT* KDMs.

### Disk diffusion assays demonstrate that MurT/GatD and NamH modulate β-lactam resistance

To validate these findings, disk diffusion assays were performed with control strains and the KDMs to assess potential susceptibility differences to β-lactams combined with clavulanate (AMX+CLA, CTX+CLA, and MEM+CLA) on solid media following CRISPRi induction (**Figure 5B**).

Based on the qRT-PCR results, only one sgRNA per target gene was selected for further assays, using two main criteria: **i)** efficient silencing of the respective target gene; **ii)** sufficient growth of the resulting KDMs in the phenotyping assays to ensure that the observed differences in susceptibility and immune response assays arise from gene knockdown rather than lethal effects, which is particularly important for essential genes. The chosen sgRNAs were sgRNA1 for the *namH* and *murT* genes and sgRNA2 for the *gatD* gene.

To evaluate susceptibility differences on solid media, antibiotics were prepared at concentrations that produced a distinguishable zone of inhibition. For all β-lactams, the employed concentrations were 256 or 512-fold higher than the MIC determined in liquid media. As anticipated, the susceptibility of *Mtb* WT and PLJR965 to β-lactams remained relatively stable, even with the inducer (**Figure 5B**). Again, the depletion of characteristic mPG modifications promoted increased susceptibility to β-lactams (**Figure 5B**). Regarding AMX+CLA, sgRNA2-mediated *gatD* repression resulted in significantly heightened susceptibility compared to the corresponding uninduced control (*P* = 0.0457). Additionally, a significant increase in susceptibility was observed upon sgRNA1-mediated *namH* silencing relative to WT ATc (*P* = 0.0196), PLJR965 ATc (*P* = 0.0142), and the respective uninduced control (*P* = 0.0457). For CTX+CLA, susceptibility was significantly increased upon *namH* silencing versus WT ATc (*P* = 0.0118) and PLJR965 ATc (*P* = 0.0253). Susceptibility to MEM+CLA was also significantly increased following sgRNA2-mediated *gatD* repression compared to the respective uninduced control (*P* = 0.0407). Likewise, sgRNA1-mediated *namH* knockdown prompted a pronounced increase in susceptibility relative to WT ATc (*P* = 0.0033) and the corresponding uninduced control (*P* = 0.0033). Generally, NamH depletion resulted in significant β-lactam hypersusceptibility. Furthermore, MEM+CLA was the most potent β-lactam against *Mtb* growth, as indicated by a broader zone of inhibition when compared to AMX+CLA and CTX+CLA (**Figure 5B**).

### NamH depletion synergizes with ethambutol and AMX/MEM+CLA

To evaluate interactions between frontline agents, β-lactams, and CRISPRi-mediated target inhibition (simulating the prospective chemical inhibition of PG modifications), checkerboard assays were performed with the control strains and selected KDMs (**Figures 6 and S3**). Given the preceding evidence of synergistic effects between ethambutol and amoxicillin-clavulanate (AMX+CLA) or meropenem-clavulanate (MEM+CLA) in *Mtb* H37Rv (48), these combinations were selected for further assessment. The fractional inhibitory concentration indices (FICIs) were calculated and the interactions between EMB and AMX/MEM+CLA were classified as synergistic, additive, indifferent, or antagonistic.

**Figure 6.**
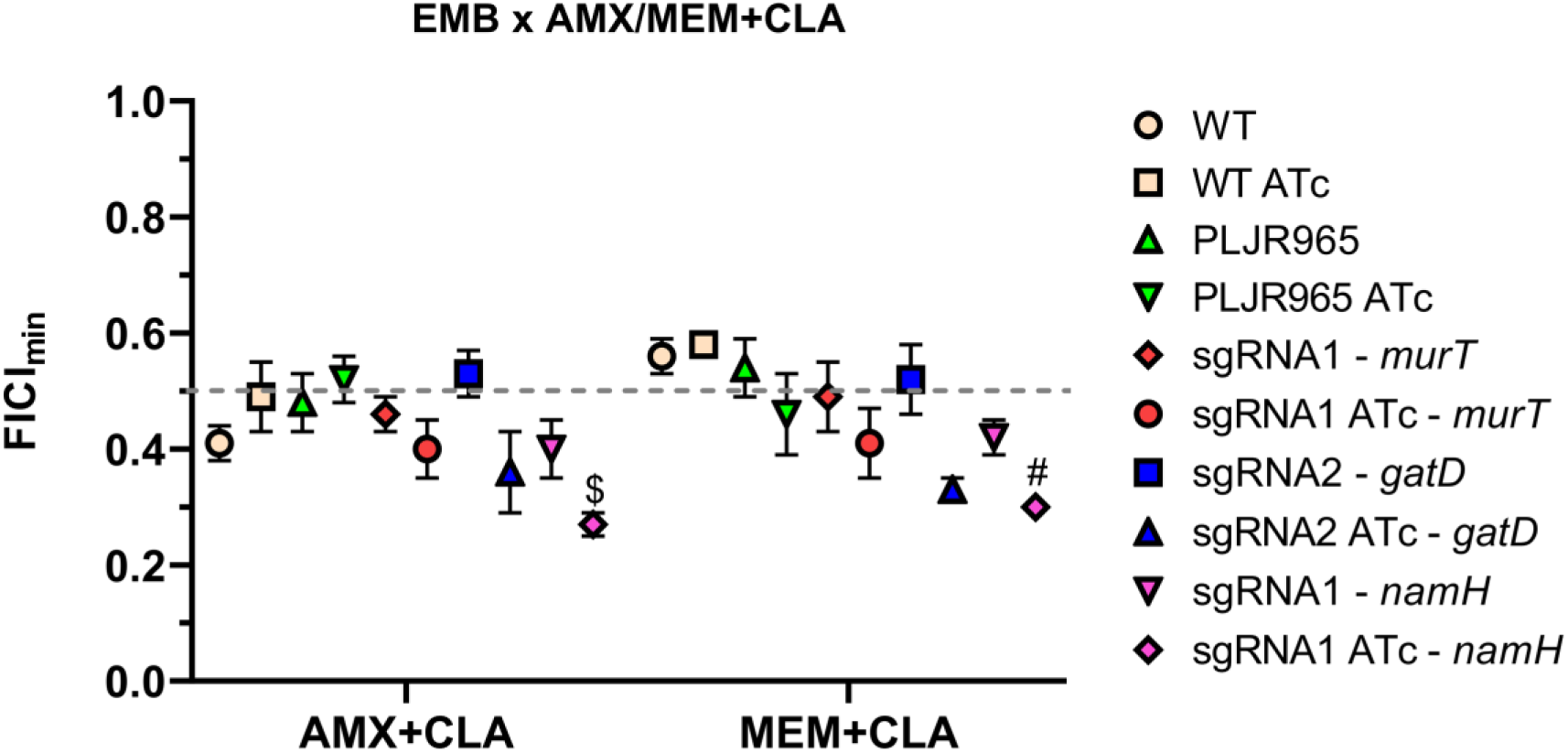
Representation of the minimum fractional inhibition indices (FICI_min_) assessing the interaction between EMB and the beta-lactams AMX+CLA and MEM+CLA were determined for the control strains (*Mtb* WT and PLJR965) and the *murT* (sgRNA1), *gatD* (sgRNA2), and *namH* (sgRNA1) knockdown mutants, with and without 100 ng/mL of ATc (n ≥ 3). The concentration of clavulanate was fixed at 5 μg/mL. AMX, amoxicillin; CLA, clavulanate; EMB, ethambutol; MEM, meropenem. Error bars show the standard error of the mean (SEM). The dotted grey line represents the FICImin value underneath which interactions are considered synergistic (FICI_min_ ≤ 0.5). Multiple comparisons were made using one-way ANOVA: * *P* < 0.05. Significant differences are indicated with symbols: # comparing to *Mtb* WT ATc and $ comparing to *Mtb* PLJR965 ATc.

The checkerboard assays showed that target repression provoked reduced minimum FICI (FICI_min_) values when assessing the interaction between EMB and AMX/MEM + CLA (**Figure 6**). All strains, except for *Mtb*::PLJR965 ATc and the uninduced *Mtb*::PLJR965-sgRNA2-*gatD*, displayed a synergistic effect between EMB and AMX+CLA. Synergies were less common with EMB plus MEM+CLA, where some interactions were classified as additive. Notably, depletion of the *N*-glycolylation of muramic acid significantly decreased the FICI_min_ between EMB plus AMX+CLA (*P* = 0.0205) or MEM+CLA (*P* = 0.0362) compared to PLJR965 ATc and WT ATc, respectively (**Figure 6**).

### D-*i*Glu amidation contributes to intracellular survival

To investigate how the distinctive mPG modifications modulate host-pathogen interactions, THP-1-derived macrophages were infected with the control strains and the KDMs at a MOI of 0.25. The intracellular survival (in CFU/mL) was evaluated at 1, 3, and 6 days post-infection.

As anticipated, the intracellular viability of *Mtb* increased over time (26, 51, 54–56). Also, the addition of the inducer had little effect on the survival of *Mtb* WT and PLJR965 within THP-1 macrophages (**Figure 7A-C**).

**Figure 7.**
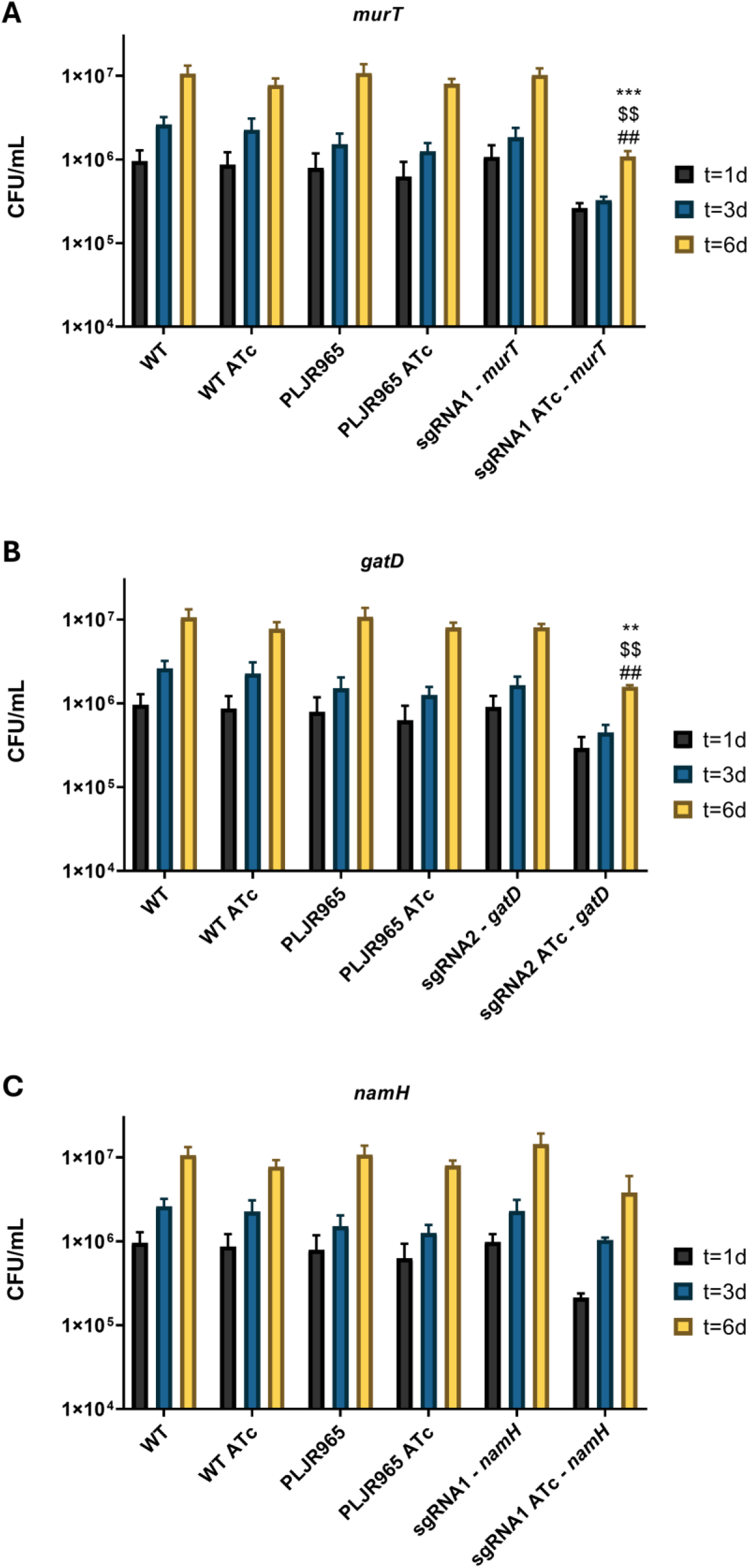
Logarithmic representation of the mean bacterial survival (in CFU/mL) after disruption of THP-1-derived macrophages infected with the control strains (*Mtb* WT and PLJR965) and with the knockdown mutants, in the presence and absence of 100 ng/mL ATc (n=3): **(A)** *murT* knockdown mutants (sgRNA1); **(B)** *gatD* knockdown mutants (sgRNA2); **(C)** *namH* knockdown mutants (sgRNA1). Error bars show the standard error of the mean (SEM). Multiple comparisons were made using one-way ANOVA: * *P* < 0.05; ** *P* < 0.01; *** *P* < 0.001. Significant differences are indicated with symbols: # comparing to *Mtb* WT ATc, $ comparing to *Mtb* PLJR965 ATc, * comparing to the respective uninduced control.

Notably, the CRISPRi-mediated knockdown of *murT*/*gatD* induced a reduction in *Mtb* intracellular survival compared to the control strains, with significant differences at 6 days post-infection. The sgRNA1-mediated knockdown of *murT* prompted a highly significant decline in intracellular survival at T_6_ relative to WT ATc (*P* = 0.0015), PLJR965 ATc (*P* = 0.0012), and the respective uninduced control (*P* = 0.0005) (**Figure 7A**). Similarly, sgRNA2-mediated *gatD* knockdown led to a very significant reduction in intracellular viability at T_6_ compared to WT ATc (*P* = 0.0054), PLJR965 ATc (*P* = 0.0039), and the respective uninduced control (*P* = 0.0027) (**Figure 7B**). Conversely, the reduction in intracellular survival caused by sgRNA1-mediated *namH* knockdown was not considered statistically significant (**Figure 7C**).

### D-*i*Glu amidation is a mechanism of immune evasion

To uncover whether these PG modifications influence the immune response upon *Mtb* infection of THP-1-derived macrophages, the cells were lysed 6 days post-infection, and the mRNA levels of the pro-inflammatory cytokines TNF-α and IL-1β and of the anti-inflammatory cytokine IL-10 were evaluated by qRT-PCR (**Figure 8**).

**Figure 8.**
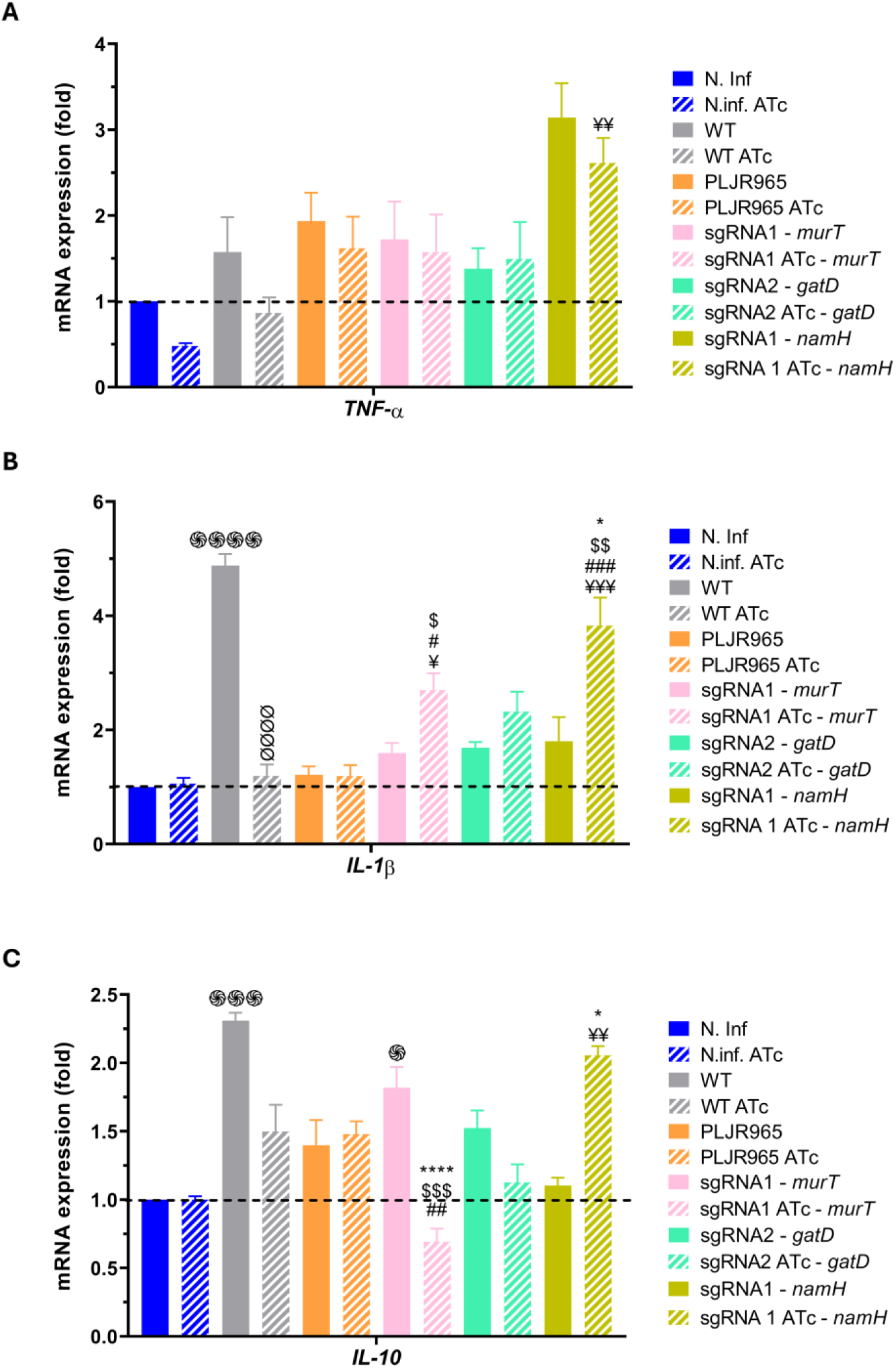
mRNA expression levels of (A) TNF-α, (B) IL-1β, (C) IL-10 assessed in THP-1-derived macrophages infected with the controls (*Mtb* WT and PLJR965) and the *murT*, *gatD*, and *namH* knockdown mutants, with and without ATc (n=3). Cytokine expression was assessed at 6 days post-infection and normalized to *GAPDH*. Smooth bars represent infections with uninduced strains, while striped bars depict ATc-induced conditions. Dashed lines represent non-infected macrophages as the calibrator. Error bars show the standard error of the mean (SEM). Multiple comparisons were made using one-way ANOVA: * *P* < 0.05; ** *P* < 0.01; *** *P* < 0.001. Significant differences are depicted with symbols: ֍ versus non-infected macrophages, ¥ vs. N. Inf ATc, Ø vs. *Mtb* WT, # vs. *Mtb* WT ATc, $ vs. *Mtb* PLJR965 ATc, * vs. the respective uninduced control.

Overall, TNF-α levels showed no major changes, although a non-significant trend suggested slightly higher expression in infected versus non-infected macrophages (N. Inf) (**Figure 8A**). Although not significant, non-infected ATc-treated macrophages displayed about half the TNF-α expression levels of their untreated counterparts. Remarkably, TNF-α levels were significantly increased upon infection with the induced *Mtb*::PLJR965-sgRNA1-*namH* KDM, compared to non-infected ATc-treated macrophages (N. Inf ATc; 5.47-fold; *P* = 0.0066) (**Figure 8A**). However, no significant differences were observed between induced and non-induced sgRNA1-*namH* strains, as both stimulated similar TNF-α expression levels (**Figure 8A**).

Furthermore, non-infected untreated and ATc-treated macrophages exhibited similar IL-1β levels. Nevertheless, infection with *Mtb* WT caused a highly significant increase in IL-1β expression (4.88-fold; *P* < 0.0001 vs. N. Inf), an effect that was absent with *Mtb* WT ATc (**Figure 8B**). Unexpectedly, infection with *Mtb*::PLJR965 reduced IL-1β levels relative to WT but did not provoke any significant expression changes compared to non-infected macrophages. The mRNA levels of IL-1β increased significantly following sgRNA1-mediated *murT* silencing compared to N. Inf ATc (2.55-fold; *P* = 0.0146), WT ATc (2.26-fold; *P* = 0.0350), and PLJR965 ATc (2.26-fold; *P* = 0.0398) (**Figure 8B**). A similar tendency was observed following the sgRNA2-mediated silencing of *gatD*, although without significance. Likewise, IL-1β expression was also significantly elevated upon sgRNA1-mediated *namH* knockdown relative to N. Inf ATc (3.61-fold; *P* = 0.0004), WT ATc (3.20-fold; *P* = 0.0010), PLJR965 ATc (3.21-fold; *P* = 0.0011), and the respective uninduced control (2.12-fold; *P* = 0.0485) (**Figure 8B**).

Infection with *Mtb* WT caused a highly significant rise in IL-10 expression relative to N. Inf (2.31-fold; *P* = 0.0004) (**Figure 8C**). Moreover, sgRNA1-mediated *murT* knockdown provoked a highly significant decrease in IL-10 levels compared to WT ATc (2.16-fold; *P* = 0.0011), PLJR965 ATc (2.14-fold; *P* = 0.0009), and the respective uninduced control (2.63-fold; *P* < 0.0001) (**Figure 8C**). A similar tendency was observed following the sgRNA2-mediated silencing of *gatD*, yet without statistical significance. Conversely, sgRNA1-mediated *namH* silencing significantly increased IL-10 levels versus N. Inf ATc (2.06-fold; *P* = 0.0025) and the respective uninduced control (1.86-fold; *P* = 0.0122) (**Figure 8C**).

## DISCUSSION

Studying PG biosynthesis enzymes and inhibitors offers a promising approach to uncover therapeutic targets essential for the development of innovative anti-TB agents against drug-resistant TB. Alongside increasing clinical reports demonstrating the benefits of β-lactam/β-lactam inhibitor (BL/BLI) combinations against MDR-TB (61–66), the WHO has included meropenem and imipenem-cilastatin in group C of drugs advised for extended MDR-TB regimens (67). Here, we investigated the roles of D-*i*Glu amidation and of the *N*-glycolylation of muramic acid in *Mtb* growth, β-lactam susceptibility, and host immune modulation. Our findings, discussed below, revealed that MurT/GatD and NamH contribute to β-lactam resistance, with MurT/GatD also playing a fundamental role in *Mtb* intracellular survival and immune evasion.

The qRT-PCR results demonstrated effective silencing of all GOIs with at least one of the employed sgRNAs. Contrasting previous observations in *Msm* (9, 46) but consistent with findings in *Mtb* (42, 43, 68), no correlation was found between target inhibition efficiency and the distance between the target site and the transcription start site. Indeed, *gatD* knockdown was more efficient when mediated by sgRNA2-*gatD* despite employing a weaker PAM and targeting a distal ORF region. Hence, factors like GC content and sgRNA length may justify this peculiar observation (68). While no leaky expression was observed, target repression was lower than what was previously described for dCas9_Sth1_-mediated targeting of selected GOIs in *Mtb* (12–99 fold), likely owing to the relatively shorter induction time utilized here (3 *vs*. 4 days) (42). Collectively, our results suggest that dCas_Sth1_-mediated knockdown produces effective target mRNA inhibition in *Mtb*, directly impacting the associated phenotypes.

After, the phenotyping assays confirmed that D-*i*Glu amidation is essential for *Mtb* survival, supporting previous observations in *Msm* (9) and *Mtb* (8, 22, 23, 59). The conserved nucleotide regions producing the first domain of MurT_TB_, Mur_Ligase_M, a known binding site of cofactors ATP/Mg^2+^, are targeted by sgRNA2-*murT* (8, 13). In this case, sgRNA2-guided *murT* repression culminates in the production of a small mRNA that is probably degraded, completely abrogating protein synthesis and ultimately causing the lethal phenotype displayed by the corresponding KDM. Within the second domain of MurT_TB_, DUF1727, sgRNA1-*murT* targets the four conserved amino acid regions (I-IV) that govern the MurT-GatD interplay (8, 9, 69). Nevertheless, since the target site is near the 3’-end of *murT* and the employed PAM (PAM9) is relatively weak, RNA polymerase may still transcribe most of *murT*, generating a semi-functional MurT protein and facilitating the growth of the respective KDM (9). Conversely, the glutamine amidotransferase (GATase_3) domain of GatD_TB_ encompasses conserved residues responsible for glutamine capture (R124) and for the channeling of ammonia to MurT_TB_ (G56) along with catalytic residues in GatD_TB_ (C90, H190) and in MurT_TB_ (D337) (5, 8, 9, 13). Repression of *gatD* with sgRNAs 1 and 2, which target sites near some of these residues, may lead to depleted D-*i*Glu amidation. Since only sgRNA2-mediated repression caused a significant *gatD* knockdown, the corresponding KDM displays impaired growth.

In contrast, *namH* is non-essential for *Msm* growth (9) and generally classified as non-essential for *Mtb* viability (22, 23, 60). Nonetheless, sgRNA2-mediated repression of *namH* resulted in discernible growth deficits on solid media, implying that NamH may impact *Mtb* fitness under certain conditions. Together with a previous study that classified *namH* as essential for *Mtb* growth (59), our findings suggest that *namH* may be conditionally essential. NamH comprises a N-terminal UlaG domain of unknown function and a C-terminal Rieske domain, responsible for the *N*-glycolylation of muramic acid (9, 20). The sgRNAs employed here target the 5’-end of *namH*, corresponding to the UlaG domain. Even though sgRNA1-*namH* targets the promoter zone, the more pronounced growth inhibition phenotype observed with sgRNA2-*namH* may stem from differences in PAM strength and/or specific target site properties.

Furthermore, the susceptibility assays demonstrated that both D-*i*Glu amidation and the *N*-glycolylation of muramic acid contribute to β-lactam resistance. In fact, NamH and MurT/GatD have been shown to modulate β-lactam resistance in mycobacteria (9, 10, 20) and *S. aureus* (11, 13). Globally, sgRNA2-mediated *murT* silencing provoked the greatest susceptibility differences relative to *Mtb* WT, suggesting that the catalytic activity of MurT plays a role in β-lactam resistance. The observation that minimal sgRNA1-mediated *gatD* silencing caused greater β-lactam susceptibility differences than significant sgRNA2-mediated silencing suggests that targeting the 5’-region of *gatD* has a more profound impact on enzyme function and CW integrity. Moreover, susceptibility gains were often correlated with growth defects in the phenotyping assays, namely in the *murT* and *namH* KDMs.

Additionally, CTX is typically more effective against *Mtb* than AMX, since it inhibits both PBPs and Ldts (71). Curiously, AMX+CLA was more potent than CTX+CLA because clavulanate significantly enhances the activity of AMX, while CTX alone is more stable against BlaC-mediated hydrolysis. Furthermore, the greater MIC reduction observed with CLA plus AMX (_∼_64-fold) versus CTX or MEM (_∼_4-fold) supports the fact that BlaC preferentially hydrolyzes penicillins over cephalosporins or carbapenems (33, 70). In accordance with previous findings in *Msm* (9), the higher antimycobacterial activity observed for MEM versus AMX and CTX is likely attributed to its resistance against BlaC-mediated hydrolysis together with its dual inhibition of both PBPs and Ldts (9, 72). Curiously, the increased susceptibility of *murT* KDMs to INH, consistent with earlier findings in *Msm* (9), suggests a role for MurT in reduced susceptibility to INH.

Collectively, the disk diffusion assays verified increased β-lactam susceptibility following the CRISPRi-mediated inhibition of characteristic mPG modifications. Unlike MurT, GatD depletion significantly increased susceptibility to AMX+CLA and MEM+CLA, consistent with a role for D-*i*Glu amidation in β-lactam resistance. Notably, the depletion of NamH provoked a highly significant increase in β-lactam susceptibility, underscoring the relevance of the *N*-glycolylation of muramic acid to the intrinsic β-lactam resistance phenotype of WT mycobacteria. Therefore, the development of antibiotics targeting these mPG modifications could synergize with β-lactams and frontline antimycobacterials to advance TB treatment. Again, MEM+CLA was the most potent β-lactam against mycobacteria, reinforcing its value as a prospective addition to combination therapy (9).

The checkerboard assays demonstrated that the inhibition of characteristic mPG modifications produced reduced FICI_min_ values between EMB and AMX/MEM + CLA. Consistent with previous findings showing a lower FICI_min_ for a combination of EMB plus AMX+CLA versus MEM+CLA in *Mtb* WT (48), synergistic effects were more commonly observed with AMX+CLA. The stronger synergy observed with EMB plus AMX+CLA might stem from the substantial enhancement of AMX efficacy when combined with clavulanate, which is remarkably superior to that observed with MEM+CLA. Also, the FICI_min_ values obtained for the EMB and MEM+CLA combination in *Mtb* WT did not meet the conventional threshold for synergy (FICI ≤ 0.5), rather indicating an additive effect, although they were roughly identical to previously reported values (48). Recently, several studies have reported synergies between penicillins, carbapenems, and cephalosporins with frontline antimycobacterials rifampicin and ethambutol against *Mtb in vitro* and intracellularly (65, 73, 74). Remarkably, orally administered cephradine and faropenem exhibit synergism with novel anti-TB agents bedaquiline, delamanid, and pretomanid (65). Here, CRISPRi-mediated NamH depletion, coupled with EMB-induced AG biosynthesis inhibition, and β-lactam-mediated transpeptidase inhibition, resulted in synergy. Together with the fact that NamH depletion causes β-lactam hypersusceptibility, these findings support the hypothesis that NamH inhibition could synergize with therapeutic schemes containing EMB and AMX/MEM+CLA.

Investigating how the targeted mPG modifications modulate interactions with the host could guide the development of novel therapeutics that strengthen immune responses against TB infection. The host pattern recognition receptors (PRRs)-mediated recognition of mPG, a conserved pathogen-associated molecular pattern (PAMP), induces pro-inflammatory responses (e.g., elevated TNF-α and IL-1β levels) against *Mtb*. Soluble PG fragments containing D-glutamyl-*meso*-diaminopimelic acid and muramyl dipeptide (MDP) are recognized by intracellular PRRs NOD1 and NOD2, respectively (75, 76).

In line with this, infection assays showed that depletion of MurT/GatD reduced *Mtb* intracellular survival, likely by unmasking NOD1 ligands and consequently stimulating macrophage activation and immune cell recruitment (77), alluding to a crucial role for D-*i*Glu amidation in persistence. These findings reflect previous *in vitro* studies in *Msm* (9) and *in vivo* studies in *M. bovis* BCG (19). Conversely, NamH activity was found to be non-essential for *Mtb* intracellular survival, supporting former studies (26). Even though we had previously demonstrated that NamH activity contributes to the survival of *Msm* within host macrophages (9), this effect was not observed with pathogenic *Mtb*.

Moreover, depletion of D-*i*Glu amidation caused upregulated IL-1β mRNA levels and downregulated IL-10 expression, while the inhibition of the *N*-glycolylation of muramic acid induced the expression of both IL-1β and IL-10.

Non-infected ATc-treated THP-1-derived macrophages displayed half the TNF-α expression levels of their untreated counterparts, suggesting that ATc may specifically modulate TNF-α expression in non-infected cells through an unknown mechanism. NamH depletion, which results in a higher proportion of *N*-acetyl-MDP, increased TNF-α expression versus non-infected ATc-treated macrophages, though this effect was not significant compared to the uninduced control. Studies in which mouse peritoneal macrophages were infected with NamH-depleted mycobacteria indicated that *N*-glycolyl-MDP is more immunogenic than *N*-acetyl-MDP at stimulating NOD2-mediated TNF-α production, as measured by enzyme-linked immunosorbent assay (ELISA) (25, 26, 78). In a previous study, there were no differences in intracellular survival, while TNF-α secretion was significantly decreased in human monocyte-derived macrophages infected with Δ*namH* mutants versus WT *Mtb* (26). By contrast, another study showed that the *N*-glycolylation of muramic acid does not significantly affect human NOD1/2-mediated immune recognition (17). Discrepancies between studies using mouse- and human-derived macrophages might be attributed to species-specific basal gene expression and responsiveness of NOD sensors. Overall, the observed differences were not statistically significant, as robust TNF-α expression is mostly elicited when NOD2 is co-activated with other PRRs (79).

On top of that, macrophage infection with *Mtb* WT significantly increased IL-1β expression relative to non-infected controls, whereas infection with *Mtb* WT ATc did not alter IL-1β mRNA levels. Despite having little effect on *Mtb* growth (80), ATc may exert immunomodulatory effects, thereby preventing macrophage activation (81). Similarly, macrophage infection with *Mtb*::PLJR965 did not affect IL-1β levels, perhaps owing to altered PAMP secretion or metabolic stress responses that dampen pro-inflammatory signaling. Notably, MurT/GatD depletion upregulated IL-1β levels, indicating heightened immune recognition. Supporting this, several reports uncovered that D-*i*Glu amidation mediates decreased NOD1/2-mediated immune recognition (16–18). Accordingly, MurT/GatD depletion in *M. bovis BCG* provoked increased NOD1/2-mediated immune recognition (19). Furthermore, NamH depletion upregulated IL-1β levels, also implying enhanced immune recognition.

Despite both being pro-inflammatory cytokines, the mRNA levels of TNF-α and IL-1β differed, likely due to distinct sensitivities to transcriptional regulation, as IL-1β expression, unlike TNF-α, is readily induced through NF-κB signalling (82).

In this context, the regulation of IL-10 levels during TB infection is crucial to prevent excessive inflammation and tissue destruction. Indeed, excessive IL-10 levels facilitate *Mtb* persistence by inhibiting pro-inflammatory cytokines, phagocytosis, and macrophage activation (83, 84). Our results demonstrated that the depletion of MurT/GatD downregulated IL-10 mRNA levels, thereby implying that D-*i*Glu amidation promotes immune evasion and mycobacterial persistence (16, 17, 19). Conspicuously, NamH depletion provoked increased IL-10 expression, likely as a compensatory response to the concurrent rise in IL-1β levels. Conversely, the *N*-glycolylation of muramic acid did not impact IL-10 expression.

Overall, our findings show that while the *N*-glycolylation of PG does not significantly impact *Mtb* intracellular survival, D-*i*Glu amidation facilitates persistence by promoting immune evasion strategies, such as the production of anti-inflammatory cytokines.

Our work has numerous limitations. First, CRISPRi is typically associated with downstream polarity in operonic genes (45, 46). Nevertheless, since these genes are often co-transcribed and functionally related, this issue is rather minimized (42, 68). Despite endorsing the development of small molecules targeting MurT/GatD and NamH, we recognize that their ability to fully suppress enzyme functions may be limited compared to CRISPRi (68). Besides, immune response differences were assessed by qRT-PCR instead of ELISA, which would have enabled direct cytokine quantification and facilitated comparison with previous studies.

To conclude, both PG modifications contribute to β-lactam resistance. While NamH is a key resistance determinant, the activity of MurT/GatD sustains intracellular survival by enabling immune evasion and limiting the efficacy of antimycobacterials. Our findings in *Mtb* suggest that targeting MurT/GatD and NamH, either individually or simultaneously, could represent a novel TB-fighting strategy, with the bonus of improving β-lactam efficacy in MDR-TB therapy.

## AUTHOR CONTRIBUTIONS

C.S., F.O., D.P., E.A. and M.J.C. conceived and designed the study. C.S., M.M., and F.O. performed the experiments. C.S., F.O., D.P., E.A. and M.J.C. analyzed the data. C.S. wrote the original draft. All authors reviewed and approved the final version of the manuscript.

## DATA AVAILABILITY STATEMENT

All data supporting the findings of this study are included in the article and supplementary materials. Additional information is available from the corresponding author upon request.

## Supplementary MATERIAL

FIG. S1 TO S3, TABLES S1-S4

