## Supplementary Material for "Genetic interference of distinctive *Mycobacterium tuberculosis* peptidoglycan modifications enhances β-lactam susceptibility and reveals expression-sensitive host immune dynamics"

### 1. Supplementary Figures and Tables

#### 1.1. Supplementary Figures

***Mtb*::PLJR965-sgRNA1 *murT* – Sequencing output**

5' – ...ATCAGTGATAGATATAATCTGGGAGCCGGCCGGGTTTTTGGCCAGTTTTTGTACTCGAAAGAAGCTACAAAGATAAGGCTTCATGCCGTAATC... - 3'

***Mtb*::PLJR965-sgRNA2 *murT* – Sequencing output**

5' - ...ATCAGTGATAGATATAATCTGGGAGTCCAGCTGGTCTCGGGAGAGGTGTTTTTGTACTCGAAAGAAGCTACAAAGATAAGGCTTCATGCCGAAATC... – 3'

***Mtb*::PLJR965-sgRNA1 *gatD* – Sequencing output**

5' – ...ATCAGTGATAGATATAATCTGGGAATCTCGGCGGCGATGCCGCGCAGTTTTTGTACTCGAAAGAAGCTACAAAGATAAGGCTTCATGCCGAAATC... – 3'

***Mtb*::PLJR965-sgRNA2 *gatD* – Sequencing output**

5' - ... ATCAGTGATAGATATAATCTGGGAGCCCAAGGGCGACGTTCCGGGCCGTTTTTGTACTCGAAAGAAGCTACAAAGATAAGGCTTCATGCCGAAATC... - 3'

***Mtb*::PLJR965-sgRNA1 *namH* – Sequencing output**

5' – ...CCTGGGGATTTGCGATCCCTATCAGTGATAGATATAATCTGGGAGTGACCTGCACAGCTACCTTGTTTTTGTACTCGAAAGAAGCTACAAAGATAAGGCTTCATGCCGAAATCAACACCCT... – 3'

***Mtb*:: PLJR965-sgRNA2 *namH* – Sequencing output**

5' – ...TGGGGATTTGCGATCCCTATCAGTGATAGATATAATCTGGGAGCTGCCGGCCTGGGTCTGGAGTTTTTGTACTCGAAAGAAGCTACAAAGATAAGGCTTCATGCCGAAAT... – 3'

**Fig. S1 -** Sequencing results (5' – 3') of PLJR965 with the sgRNA targeting each gene of interest (*murT, gatD*, or *namH*) in *M. tuberculosis*. The -10 site is underlined in yellow, the insert or sgRNA is underlined in green and the dCas9 handle is painted in grey.

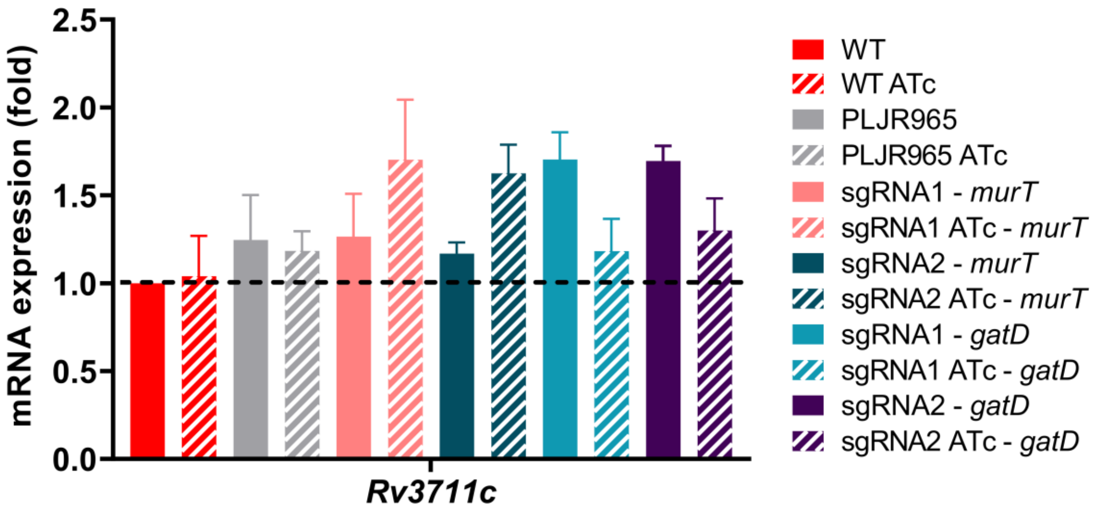


**Fig. S2** - Mean of the relative mRNA expression levels of *Rv3711c* (gene immediately upstream of *murT*), normalized to *sigA*, at 72 hours post-induction, with (stripped bars) and without (smooth bars) 100 ng/mL ATc (n=3): *murT* and *gatD* knockdown. The dashed lines show the WT sample as calibrator. Error bars show the standard error of the mean (SEM). Multiple comparisons were made using one-way ANOVA: * $P < 0.05$; ** $P < 0.01$; *** $P < 0.001$. Significant differences are indicated with symbols: # compared to *Mtb* WT ATc, $ compared to *Mtb* PLJR965 ATc, * versus the respective uninduced control. The • symbol indicates a highly significant difference ($P < 0.0001$) from all controls.

| *M. tuberculosis* | | Average | | | STD DEV | | | N | S.E.M. | | |
|---|---|---|---|---|---|---|---|---|---|---|---|
| | | FICImin | FICImed | FICImax | | | | | | | |
| AMX+CLA x EMB | WT | 0.41 | 0.57 | 0.95 | 0.05 | 0.01 | 0.19 | 4 | 0.03 | 0.01 | 0.09 |
| | WT ATc | 0.49 | 0.62 | 1.08 | 0.11 | 0.09 | 0.03 | 4 | 0.06 | 0.04 | 0.02 |
| | PLJR965 | 0.48 | 0.77 | 1.12 | 0.11 | 0.17 | 0.07 | 6 | 0.05 | 0.07 | 0.03 |
| | PLJR965 ATc | 0.52 | 0.71 | 1.02 | 0.09 | 0.10 | 0.18 | 6 | 0.04 | 0.04 | 0.07 |
| | sgRNA1 - *murT* | 0.46 | 0.78 | 0.94 | 0.06 | 0.18 | 0.27 | 3 | 0.03 | 0.10 | 0.16 |
| | sgRNA1 ATc - *murT* | 0.40 | 0.58 | 1.06 | 0.08 | 0.04 | 0.00 | 3 | 0.05 | 0.02 | 0.00 |
| | sgRNA2 - *gatD* | 0.53 | 0.72 | 1.11 | 0.09 | 0.07 | 0.03 | 4 | 0.04 | 0.03 | 0.02 |
| | sgRNA2 ATc - *gatD* | 0.36 | 0.68 | 1.07 | 0.13 | 0.16 | 0.04 | 3 | 0.07 | 0.10 | 0.02 |
| | sgRNA1 - *namH* | 0.40 | 0.63 | 1.11 | 0.08 | 0.17 | 0.03 | 3 | 0.05 | 0.10 | 0.02 |
| | sgRNA1 ATc - *namH* | 0.27 | 0.39 | 1.05 | 0.03 | 0.05 | 0.01 | 3 | 0.02 | 0.03 | 0.01 |
| MEM+CLA x EMB | WT | 0.56 | 0.73 | 1.13 | 0.05 | 0.03 | 0.00 | 3 | 0.03 | 0.02 | 0.00 |
| | WT ATc | 0.58 | 0.69 | 1.08 | 0.03 | 0.05 | 0.03 | 3 | 0.02 | 0.03 | 0.02 |
| | PLJR965 | 0.54 | 0.70 | 0.97 | 0.10 | 0.10 | 0.24 | 4 | 0.05 | 0.05 | 0.12 |
| | PLJR965 ATc | 0.46 | 0.74 | 0.98 | 0.12 | 0.21 | 0.26 | 3 | 0.07 | 0.12 | 0.15 |
| | sgRNA1 - *murT* | 0.49 | 0.59 | 0.97 | 0.11 | 0.13 | 0.20 | 4 | 0.06 | 0.07 | 0.10 |
| | sgRNA1 ATc - *murT* | 0.41 | 0.63 | 1.02 | 0.12 | 0.14 | 0.27 | 4 | 0.06 | 0.07 | 0.13 |
| | sgRNA2 - *gatD* | 0.52 | 0.65 | 0.94 | 0.11 | 0.14 | 0.27 | 3 | 0.06 | 0.08 | 0.16 |
| | sgRNA2 ATc - *gatD* | 0.33 | 0.57 | 1.11 | 0.03 | 0.05 | 0.03 | 3 | 0.02 | 0.03 | 0.02 |
| | sgRNA1 - *namH* | 0.42 | 0.61 | 0.94 | 0.06 | 0.10 | 0.22 | 3 | 0.03 | 0.06 | 0.13 |
| | sgRNA1 ATc - *namH* | 0.30 | 0.46 | 1.05 | 0.01 | 0.06 | 0.01 | 3 | 0.01 | 0.03 | 0.01 |

Synergy (FICI < 0.5) Additive Effect (0.5 < FICI < 1) Indifferent (1 < FICI < 4)

**Fig. S3 -** Heatmap of the detailed outputs of the checkerboard assays assessing the interaction between EMB and the beta-lactams AMX+CLA and MEM+CLA for the control strains (*Mtb* WT and PLJR965) and the *murT* (sgRNA1), *gatD* (sgRNA2) and *namH* (sgRNA1) knockdown mutants, with and without 100 ng/mL of ATc (n ≥ 3). The results show the values of lowest fractional inhibitory concentration (FIC) index ($FICI_{min}$), the median FICI ($FICI_{med}$), and the highest FICI ($FICI_{max}$) calculated as the average of at least three independent replicates for each combination and each strain. The color of the cells depicts the type of interaction according to the FICI value. AMX, amoxicillin; CLA, clavulanate; EMB, ethambutol; MEM, meropenem.

#### 1.2. Supplementary Tables

**Table S1 – sgRNAs used to target *murT*, *gatD*, and *namH* in *M. tuberculosis*.** [a] - PAM strength was assessed in accordance with the table repertoire of functional PAMs *in vivo* for $dCas9_{Sth1}$ defined by Rock et al., 2017 (42); [b] - mRNA knockdown caused by the electroporation of the indicated sgRNA in the *M. tuberculosis* WT strain was measured by qRT-PCR after ATc induction. Fold knockdown efficiency is relative to an uninduced control plasmid containing the same sgRNA.

| Target gene | Name of the oligos | PAM | PAM strength[a] | Gene Location 5'-3' (bp) | Base-pairing region of the sgRNA sequence (the 12 bp seed region is underlined) | BLASTed sequence in the genome of *Mtb* (seed region + PAM) | Fold knockdown efficiency[b] |
|---|---|---|---|---|---|---|---|
| ***murT*** (*Rv3712*) | sgRNA1 | 5' - GCAGGAT - 3' | PAM9 | 927 | 5' - GCCGGCCGGGTTTTTGGCCA - 3' | 5' - GGTTTTTGGCCAGCAGGAT - 3' | 10.1 |
| | sgRNA2 | 5' - TGAGCAA - 3' | PAM10 | 411 | 5' - GTCCAGCTGGTCTCGGGAGAGGT - 3' | 5' - CTCGGGAGAGGTTGAGCAA - 3' | 5.7 |
| ***gatD*** (*Rv3713*) | sgRNA1 | 5' - GCAGCAG - 3' | PAM13 | 110 | 5' - ATCTCGGCGGCGATGCCGCGCA - 3' | 5' - CGATGCCGCGCAGCAGCAG - 3' | 1.9 |
| | sgRNA2 | 5' - CGAGGAC - 3' | PAM15 | 486 | 5' - GCCCAAGGGCGACGTTCCGGGCC - 3' | 5' - ACGTTCCGGGCCCGAGGAC - 3' | 7.0 |
| ***namH*** (*Rv3818*) | sgRNA1 | 5' - TCGGAAG - 3' | PAM4 | 11 | 5' - GTGACCTGCACAGCTACCTT - 3' | 5' - CACAGCTACCTTTCGGAAG - 3' | 8.3 |
| | sgRNA2 | 5' - TCAGAAA - 3' | PAM3 | 57 | 5' - GCTGCCGGCCTGGGTCTGGA - 3' | 5' - CCTGGGTCTGGATCAGAAA - 3' | 8.3 |

**Table S2 - Primers designed and synthesized to clone the targeting sgRNAs into PLJR965 for CRISPRi-mediated targeting in *M. tuberculosis*.** [a] – PAM strength was described in accordance with the table of functional PAMs *in vivo* for dCas9$_{Sth1}$-mediated targeting in mycobacteria defined by Rock *et al.,* 2017 (42); [b] – Calculated with the Eurofins Genomics Melting temperature (Tm) formula.

| Target gene | Name of the oligos | PAM | PAM strength[a] | Gene Location 5'-3' (bp) | Primers | | Length (bp) | % GC | Tm[b] (° C) |
|---|---|---|---|---|---|---|---|---|---|
| ***murT*** (*Rv3712*) | sgRNA1 | 5' - GCAGGAT - 3' | PAM9 | 927 | PFwd | 5' - GGGAGCCGGCCGGGTTTTTGGCCA - 3' | 24 | 71 | 71 |
| | | | | | PRv | 5' - AAACTGGCCAAAAACCCGGCCGGC - 3' | | 63 | 68 |
| | sgRNA2 | 5' - TGAGCAA - 3' | PAM10 | 411 | PFwd | 5' - GGGAGTCCAGCTGGTCTCGGGAGAGGT - 3' | 27 | 67 | 73 |
| | | | | | PRv | 5' - AAACACCTCTCCCGAGACCAGCTGGAC - 3' | | 59 | 70 |
| ***gatD*** (*Rv3713*) | sgRNA1 | 5' - GCAGCAG - 3' | PAM13 | 110 | PFwd | 5' - GGGAATCTCGGCGGCGATGCCGCGCA - 3' | 26 | 73 | 74 |
| | | | | | PRv | 5' - AAACTGCGCGGCATCGCCGCCGAGAT - 3' | | 65 | 71 |
| | sgRNA2 | 5' - CGAGGAC - 3' | PAM15 | 486 | PFwd | 5' - GGGAGCCCAAGGGCGACGTTCCGGGCC - 3' | 27 | 78 | 77 |
| | | | | | PRv | 5' - AAACGGCCCGGAACGTCGCCCTTGGGC - 3' | | 70 | 74 |
| ***namH*** (*Rv3818*) | sgRNA1 | 5' - TCGGAAG - 3' | PAM4 | 11 | PFwd | 5' - GGGAGTGACCTGCACAGCTACCTT - 3' | 24 | 58 | 66 |
| | | | | | PRv | 5' - AAACAAGGTAGCTGTGCAGGTCAC - 3' | | 50 | 63 |
| | sgRNA2 | 5' - TCAGAAA - 3' | PAM3 | 57 | PFwd | 5' - GGGAGCTGCCGGCCTGGGTCTGGA - 3' | 24 | 75 | 73 |
| | | | | | PRv | 5' - AAACTCCAGACCCAGGCCGGCAGC - 3' | | 67 | 70 |

**Table S3 - Primers used to quantify the mRNA expression levels of target genes *murT*, *gatD,* and *namH* by qRT-PCR in *M. tuberculosis*.**
[a] - qRT-PCR primers were designed to avoid primer secondary structures and sequence repeats, have at least 50% of GC content, and have a Tm between 57-63°C; [b] - Calculated with the Eurofins Genomics Tm formula; [c] – AE designates the amplification efficiency calculated for each pair of primers in a single-run qPCR amplification of the calibrator sample (*M. tuberculosis* WT), performed in triplicates.

| qRT-PCR primers[a] | | | | | | | |
|---|---|---|---|---|---|---|---|
| **Target gene** | | **Primer sequences** | **Length (bp)** | **% GC** | **Tm[b] (°C)** | **Product size (bp)** | **AE[c] (%)** |
| ***sigA* (*Rv2703*)** | PFwd | 5' - CGCGACATGATGTGGATCTG - 3' | 20 | 55 | 59 | 186 | 80 |
| | PRv | 5' - CCCCTTGGTGTAGTCGAACT - 3' | 20 | 55 | 59 | | |
| ***Rv3711c*** | PFwd | 5’ – GCTGCCCTAGAGAGTGC – 3’ | 17 | 65 | 58 | 95 | 87 |
| | PRv | 5’ – TCGTCGTGAGTCACCCG – 3’ | 17 | 65 | 58 | | |
| ***murT* (*Rv3712*)** | PFwd | 5’ – GGGCGGGTTCCTGACG – 3’ | 16 | 75 | 60 | 180 | 104 |
| | PRv | 5'- TGGGCATGAGGCGATGG – 3’ | 17 | 65 | 58 | | |
| ***gatD* (*Rv3713*)** | PFwd | 5’ – GGCGACGGTTTTGATGGC – 3’ | 18 | 61 | 58 | 146 | 99 |
| | PRv | 5’ – TCCACCTCGGGCAAATCC – 3’ | 18 | 61 | 58 | | |
| ***namH* (*Rv3818*)** | PFwd | 5’ – GTGTTCCTGGATCAGATGCG – 3’ | 20 | 55 | 59 | 149 | 92 |
| | PRv | 5’ – TTGTCGGTGGTAAAGATGGC – 3’ | 20 | 50 | 57 | | |
| ***Rv3819*** | PFwd | 5' - TGGCTGCCACTGGACGTG - 3' | 18 | 67 | 61 | 142 | 85 |
| | PRv | 5' - GGTGGAGGGCGAGCAACTC - 3' | 20 | 68 | 63 | | |

**Table S4 - Primers used to quantify the mRNA expression levels of target genes *TNF-α*, *IL-1β*, and *IL-10* by qRT-PCR in *Mtb*-infected THP-1-derived macrophages.** [a] - qRT-PCR primers were designed to avoid primer secondary structures and sequence repeats, have at least 50% of GC content, and have a Tm between 57-63°C; [b] - Calculated with the Eurofins Genomics Tm formula; [c] – AE designates the amplification efficiency calculated for each pair of primers in a single-run qPCR amplification of the *Mtb* PLJR965-infected THP-1-macrophage sample, performed in triplicates.

| qRT-PCR primers[a] | | | | | | | | |
|---|---|---|---|---|---|---|---|---|
| **Target transcript** | | **Primer sequences** | **Length (bp)** | **% GC** | **Tm[b] (°C)** | **Product size (bp)** | **AE[c] (%)** | **Reference** |
| GAPDH | PFwd | 5' - CTGGGCTACACTGAGCACC - 3' | 19 | 63 | 61 | 101 | 86 | Jiang & Li, 2022 |
| | PRv | 5' - AAGTGGTCGTTGAGGGCAATG - 3' | 21 | 52 | 60 | | | |
| TNF-α | PFwd | 5' - CCAGGGACCTCTCTCTAATC - 3’ | 20 | 55 | 59 | 83 | 112 | Shu et al., 2022 |
| | PRv | 5' - ATGGGCTACAGGCTTGTCACT - 3’ | 21 | 52 | 60 | | | |
| IL-1β | PFwd | 5' - TACTCACTTAAAGCCCGCCT - 3' | 20 | 50 | 57 | 77 | 94 | Medeiros-Furquim et al., 2022 |
| | PRv | 5' - ATGTGGGAGCGAATGACAGA - 3' | | | | | | |
| IL-10 | PFwd | 5' - GCTGGAGGACTTTAAGGGTTACCT - 3' | 24 | 50 | 63 | 109 | 117 | Demers-Mathieu, Huston and Dallas, 2020 |
| | PRv | 5' - CTTGATGTCTGGGTCTTGGTTCT - 3' | 23 | 48 | 61 | | | |
